# MYRF is Essential in Mesothelial Cells to Promote Lung Development and Maturation

**DOI:** 10.1101/2025.02.13.635155

**Authors:** Gidsela Luna, Jamie Verheyden, Chunting Tan, Estelle Kim, Michelle Hwa, Jugraj Sahi, Yufeng Shen, Wendy Chung, David McCulley, Xin Sun

## Abstract

The mesothelium is a squamous monolayer that ensheathes internal organs, lines the body cavity, and the diaphragm. It serves as a protective barrier, coated in glycocalyx, and secretes lubricants to facilitate tissue movement. How the mesothelium forms is poorly understood. Here, we investigate *Myrf*, a transcription factor gene expressed in the mesothelium, because it carries variants in patients with Congenital Diaphragmatic Hernia (CDH), a disorder that affects the diaphragm, lung, and other organs. In mice, inactivation of *Myrf* early in organogenesis resulted in CDH and defective mesothelial specification, compromising its function as a signaling center for lung growth. Inactivation of *Myrf* later led to enhanced mesothelium differentiation into mesenchymal cell types through partial epithelial-to-mesenchymal transition (EMT), resulting in a unique accumulation of smooth muscle encasing the lung. In this role, MYRF functions in parallel with YAP/TAZ. Together, these findings establish MYRF as a critical regulator of mesothelium development, and when mutated, causes CDH.

## Introduction

The development of our internal organs is coordinated by complex interactions between multiple cell layers. Compared to the epithelium, mesenchyme and the endothelium, the mesothelium is poorly understood, despite its omnipresence. It is a mesoderm derived squamous epithelial monolayer that encases most of our internal organs, and also lines the body cavities and diaphragm^1^. In the adult, its canonical function is to serve as a barrier and secretes lubricants that adhere to the glycocalyx to prevent friction^2,3^. In development, the mesothelium has been shown to be important for the organogenesis of the heart, liver, and lung^4^. For example, during lung development, the mesothelium is a source of signals, such as FGF9 and retinoic acid, that promote growth and branching morphogenesis^5–8^. Interestingly, mesothelial cells themselves also have progenitor properties. Cell lineage tracing studies have shown that during heart development, the epicardium, mesothelium of the heart, can undergo epithelial-mesenchymal transition (EMT) to give rise to much of the vascular smooth muscle^9^. In the lung, while the mesothelium can give rise to vascular smooth muscle cells, pericytes, and other internal cell types, it is only a minor contributor to these cell types^10–14^.

Much of the existing knowledge on the molecular control of mesothelium development came from studies of the epicardium. Epicardial cells undergo stepwise differentiation from embryonic day E11.5 to E16.5, a process regulated by the *Wt1* transcription factor gene^15^. *Wt1* is required for epicardial cell flattening, EMT, and the expression of signaling molecules^15,16^. Epicardial EMT is also modulated by the Hippo signaling pathway which is best known for its control of organ size^17^. Notably, the Hippo pathway transcription factors *Yap/Taz* serve as upstream regulators of *Wt1*, and their inactivation disrupts epicardial EMT^18^. Conversely, the *Lats 1/2* kinases restrict YAP activity and their inactivation led to increased epicardial differentiation into cardiac fibroblasts^19^. In the lung, inactivation of *Wt1* led to reduced signaling but does not impair EMT^14,20^. Additionally, HH signaling, which is received in the embryonic lung by the mesothelium among other cells, has been shown to be required for mesothelial EMT in the lung, but not in the epicardium^12^. These studies suggest that both shared and organ-specific mesothelium gene regulatory programs are at play during development.

Malformation of the mesothelium, a critical component of the diaphragm, is at the center of the pathogenesis of congenital diaphragmatic hernia (CDH). CDH is relatively common among congenital anomalies, with a birth prevalence of 1/3000 and a mortality rate of 30-40% at birth^21^. While the hallmark of CDH is herniation of the abdominal organs through a malformed diaphragm into the chest during gestation, significant mortality at birth and long-term morbidity are primarily due to respiratory failure and pulmonary hypertension. It was previously thought that lung defects in CDH were secondary to the compression caused by herniated abdominal organs. However, using tissue-specific mouse models, multiple lines of evidence from us and others show that disruption of CDH genes in the lung, but not in the diaphragm, can alter lung development independent of herniation and mechanical compression, suggesting an intrinsic requirement for these genes in lung formation and function^22,23,24^.

From human genetic studies, rare variants in over 30 genes have been robustly associated with CDH, and other variants in at least 300 other genes have also been implicated, suggesting that the genetic cause of CDH is heterogeneous^21,25–30^. From these studies, Myelin Regulatory Factor (*MYRF*) emerged as a top CDH candidate gene, with variants found in multiple independent cases of CDH^27,31^. MYRF was first identified as a transcription factor essential for the development of oligodendrocytes in the central nervous system^32^. However, its function outside of the nervous system is poorly understood, and its role in the lung is unknown.

Here, we show in the developing mouse embryo, *Myrf* is widely expressed in the mesothelium of multiple organs, as well as the pleural cavity and the diaphragm. Inactivation of *Myrf* prior to the start of lung development resulted in highly penetrant malformation of the diaphragm and organ herniation phenotypes, demonstrating that disruption of MYRF function can lead to CDH. Further analysis showed that *Myrf* is required for the specification of the lung mesothelium. We also performed inactivation at the branching stage of lung development and found that *Myrf* is required to restrict the progenitor cell properties of the lung mesothelium and to promote the ability of the differentiated progeny to delaminate and migrate away from the lung mesothelium. Disruption of these roles led to a unique accumulation of smooth muscle cells at the lung surface. This phenotype persists into adulthood, with the ectopic smooth muscle cells producing elastin. Based solely on the histological evidence, such defects mimic the presentation of Pleuroparenchymal Fibroelastosis (PPFE), a rare adult lung condition that is poorly understood^33,34^. Our results highlight the vital roles of the lung mesothelium in normal development and CDH pathogenesis, and places *Myrf* as an essential transcriptional regulator of the lung mesothelium.

## Results

### Inactivation of *Myrf* led to CDH phenotypes

To evaluate the function of *Myrf* in lung development and CDH, we first analyzed its expression pattern in the developing embryo. At embryonic day E14.5, by RNAscope, *Myrf* transcripts were detected in the mesothelium outlining various organs and body cavities, including the lung and diaphragm (Figure S1A). To address if mutation in *Myrf* could lead to CDH, we inactivated *Myrf* in the lung mesothelium and mesenchyme by mating the *Myrf^Fl/Fl^* conditional strain to the doxycycline inducible *Tbx4-rtTA/+; tetO-cre/+* strains to generate *Tbx4-rtTA/+;tetO-cre/+;Myrf^Fl/Fl^* conditional knockout mutants (hereafter *Tbx4tetOcre;Myrf*)^32,35^. Administration of a single doxycycline injection at E6.5, followed by placement on a continuous doxycycline diet, initiates cre activity in the diaphragm, lung mesothelium, and mesenchyme at the start of lung development (hereafter *Tbx4tetOcre;MyrfE6* mutant) (Figure 1A). qRT-PCR quantification of the *Myrf* transcripts showed a reduction at the site of recombination at exons 7- 8, as well as at exons within the N-terminal or C-terminal regions, likely due to nonsense-mediated decay, confirming inactivation of *Myrf* in the lung mesenchyme (Figure 1B).The remaining signal in qRT-PCR in the *Tbx4tetOcre;MyrfE6* mutant is likely due to its expression in the embryonic epithelium as detected by RNAscope (Figure S1A).

**Figure 1:**
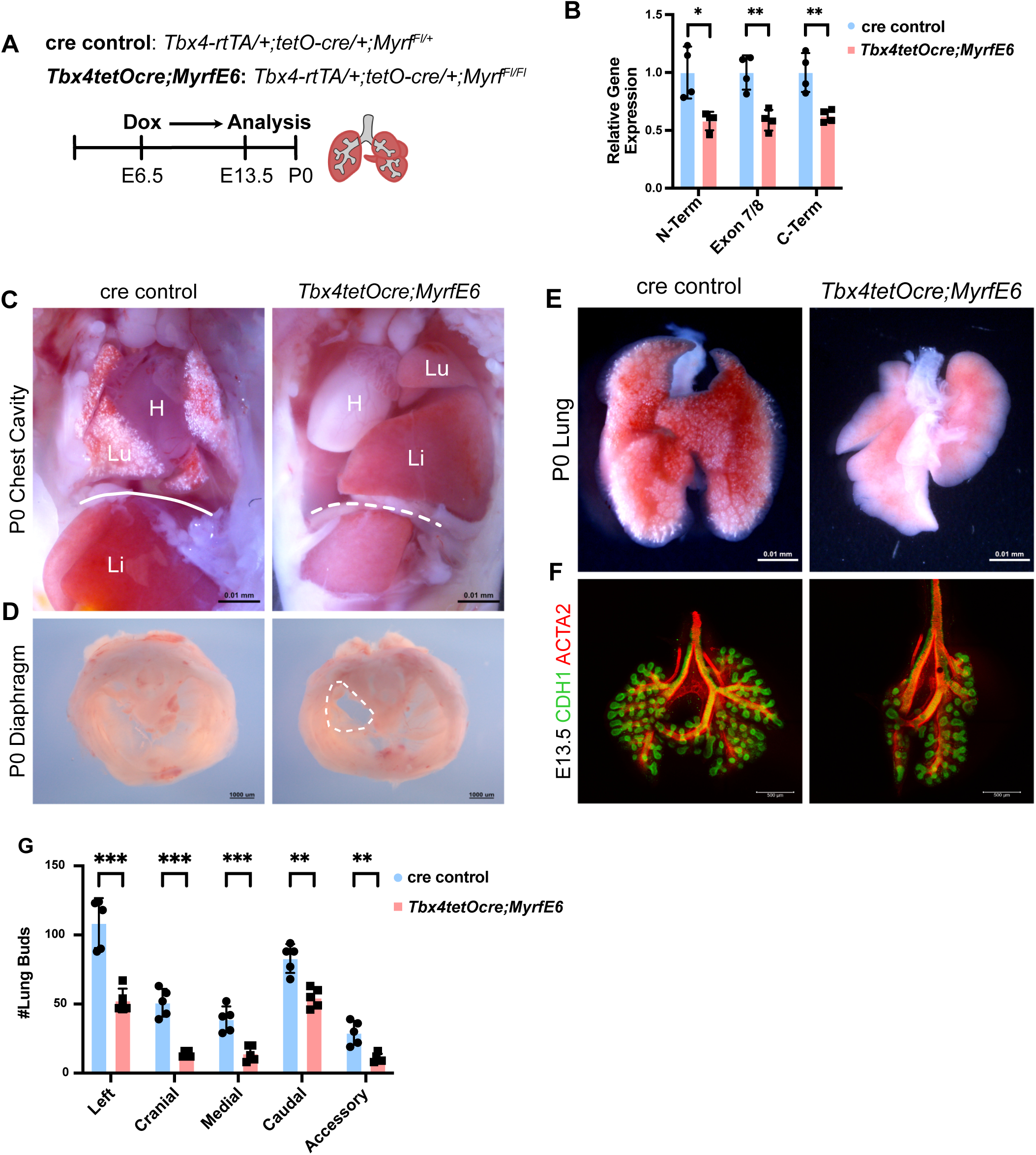
*Myrf* Promotes Lung and Diaphragm Organogenesis. **(A)** Timeline of inactivation and analysis of the *Tbx4tetOcre;MyrfE6* mutant and cre control. **(B)** qRT-PCR quantification of the *Myrf* transcript in the N-Terminus, Exon 7/8 junction, and C-Terminus regions in *Tbx4tetOcre;MyrfE6* mutant and cre control lungs. **(C)** P0 *Tbx4tetOcre;MyrfE6* mutant and cre control chest cavity. **(D)** P0 *Tbx4tetOcre;MyrfE6* and cre control diaphragm. **(E)** P0 *Tbx4tetOcre;MyrfE6* and cre control lung. **(F)** ACTA2(red) and E-Cadherin(green) whole lung immunostaining of the *Tbx4tetOcre;MyrfE6* mutant and cre control. **(G)** Lung bud tip (CDH1+) quantification in *Tbx4tetOcre;MyrfE6* mutants and cre controls at E13.5. Students T-Test was used for 1B and 1G. ns (not significant) for P ≥ 0.05, * for P < 0.05, ** for P < 0.01, *** for P < 0.001, and **** for P < 0.0001.

All *Tbx4tetOcre;MyrfE6* mutants died shortly after birth and displayed signature CDH with herniation of the abdominal contents into the chest cavity (Figure 1C). The herniation occurred on the left side of the diaphragm (n=4/4), consistent with reports that left sided CDH accounts for more than 80% of CDH cases in human (Figure 1D)^36^. The lungs of the *Tbx4tetOcre;MyrfE6* mutants were hypoplastic, misshapen, and not inflated with air at birth (Figure 1E). This inability to carry out gas exchange is likely the cause of death in the *Tbx4tetOcre;MyrfE6* mutants.

The diaphragm is derived from pleuroperitoneal folds (PPFs) which are transient embryonic structures composed of several cell types including mesothelial cells, fibroblasts, and skeletal muscles ^37,38^. Given that in the diaphragm, *Tbx4-rtTA;tetO-cre* is active both broadly in the thoracic surface of the mesothelium and in a small number of fibroblasts, we next sought to pinpoint where in the diaphragm *Myrf* is required. To determine which cell layer is contributing to the CDH phenotype, we utilized the *Prx1-cre,* which has activity in the PPF fibroblasts but not in the diaphragm mesothelium^39^. Thus, we generated *Prx1-cre;Myrf^Fl/Fl^* and *Prx1-cre;Myrf^Fl/Δ(del)^* mutants (*Prx1-cre;Myrf^Fl/Δ^* mutants carry a germline null allele of *Myrf*), which were viable and aged normally with no diaphragm or lung abnormalities (Figure S1B-D). This lack of defect, contrasting the strong phenotype in the *Tbx4tetOcre;MyrfE6* mutant, suggests that *Myrf* may be required in the mesothelium of the diaphragm for its development. These findings together demonstrate for the first time that disruption of *Myrf* function can cause CDH.

To determine if inactivation of *Myrf* reduces lung size independently of compression by herniated abdominal organs, we analyzed *Tbx4tetOcre;MyrfE6* mutant lungs at E13.5. This stage of development is prior to the closure of the diaphragm and, thus, prior to herniation. The *Tbx4tetOcre;MyrfE6* mutant displayed a hypoplastic lung with reduced number of CDH1+ lung buds across all lung lobes (Figure 1F-G,S1E-G). These data suggest that *Myrf* is required intrinsically in the lung mesenchyme for lung growth.

*Myrf* has previously been identified as an alveolar type 1 (AT1) cell marker in the adult lung^40^; however, its function in the epithelium or AT1 cells has not been addressed. We generated *Shh^cre/+^;Myrf^Fl/Fl^* (hereafter *Shhcre;Myrf*) mutants. These mutants are viable, and the lungs show normal histology at postnatal day P39, the end of lung maturation, as well as aged out to 1-year (Figure S1H-I). Mean liner intercept (MLI) was quantified at P39 and revealed no differences between the *Shhcre;Myrf* mutant and cre;het control (Figure S1L). qRT-PCR quantification for *Myrf* transcripts confirmed inactivation, while the expression of AT1 and AT2 markers (*Sftpc, Hopx, Pdpn, Aqp5* and *Ager*) remained unchanged (Figure S1J). This is confirmed by antibody staining, where the *Shhcre;Myrf* mutant displayed normal expression of HOPX and SPC and AT1:AT2 ratio (Figure S1K,M). These findings suggest the *Myrf* is not required in either epithelial cells or AT1 cells for development and homeostasis.

### Inactivation of *Myrf* compromised lung mesothelium function as a signaling center

To investigate downstream transcripts that are dysregulated in the *Tbx4tetOcre;MyrfE6* mutant lung, we performed bulk-RNAseq at E13.5, prior to closure of the diaphragm (n=4 samples for each of control and mutant). Differential gene expression analysis was performed using DESeq2 and 339 differentially expressed genes were identified (TableS1). Multiple signals for lung growth were altered in the mutant, including ones emanating from the mesothelium, as well as downstream members of these signaling pathways (Fig. 2A, TableS1). For example, members of the *Fgf9* and three FGF regulated genes, *Spry2, Etv4*, and *Etv5*, were downregulated in the *Tbx4tetOcre;MyrfE6* mutant. In contrast to *Fgf9*, which is expressed in the mesothelium aside from its expression in the epithelium, *Fgf10*, another FGF critical for lung growth, was not altered, possibly due to its broad expression in the mesenchyme (Figure 2A). *Fgf9* downregulation is especially relevant, as the “slim” lung hypoplasia phenotype of our *Tbx4tetOcre;MyrfE6* mutant closely resembles the phenotype of the *Fgf9* mutant lung^5^. The top canonical WNT gene *Wnt2* and WNT-regulated transcription effector *Lef1* were downregulated, while *Nog*, a negative regulator of BMP signaling, was upregulated. *Adlh1a2* (also known as *Raldh2*) and *Rdh10*, key enzymes involved in retinoic acid synthesis, were downregulated. Retinoic acid is a potent morphogen emanating from the mesothelium that is critical for lung development^6–8^. qRT-PCR validated the downregulation of several of these key signaling genes (Figure 2B). Together, these data demonstrate that inactivation of *Myrf* led to the dysregulation of mesothelial-derived signals essential for lung growth.

**Figure 2:**
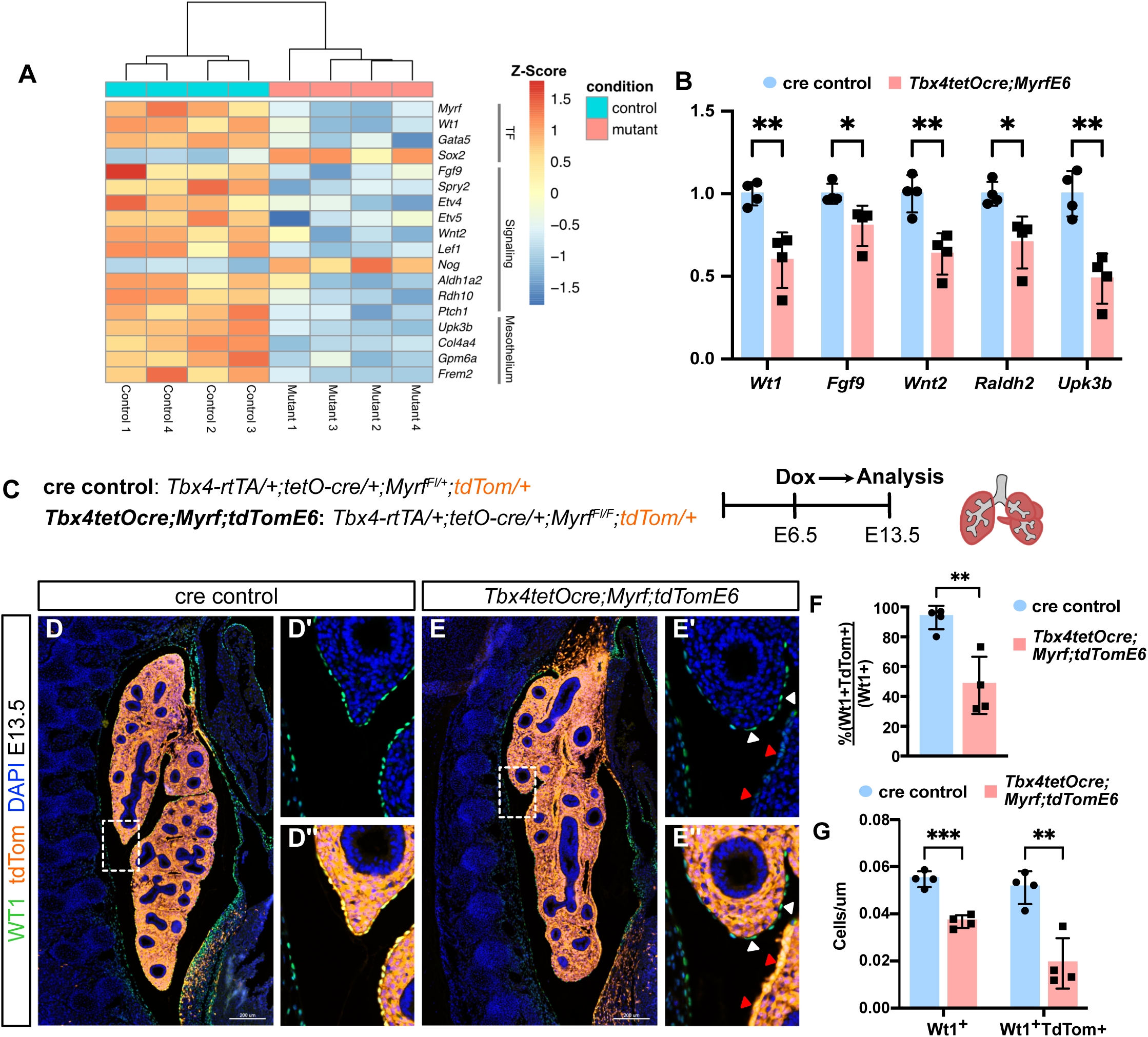
MYRF is Required for Mesothelium Specification. **(A)** Bulk RNA-Seq Heatmap of differentially expressed genes in E13.5 *Tbx4tetOcre;MyrfE6* mutants and cre controls. N=4. **(B)** qRT-PCR validation for select markers (*Wt1,Fgf9*,*Wnt2,Raldh2,* and *Upk3b).* **(C)**Timeline of inactivation and analysis of *Tbx4tetOcre;Myrf;tdTomE6* mutants and cre controls crossed to a tdTom reporter. **(D)** Immunostaining for WT1(green) and tdTom(red) in the E13.5 chest cavity of the *Tbx4tetOcre;Myrf;tdTomE6* mutant. D’ and D’’ are magnifications of D. **(E)** Immunostaining for WT1(green) and tdTom(red) in the E13.5 chest cavity of the cre control. E’ and E’’ are magnifications of E. **(F)** Percentage of WT1+TdTom+ mesothelial cells in the *Tbx4tetOcre;Myrf;tdTomE6* mutant and cre control. Wt1+ = Wt1^hi^ + Wt1^low^. **(G)** Quantification of WT1+ and WT1+tdTom+ cells in the *Tbx4tetOcre;Myrf;tdTomE6* mutant and cre control normalized to the perimeter of the lung. Wt1+ = Wt1^hi^ + Wt1^low^. Students T-Test was used for 2B, 2F and 2G. ns (not significant) for P ≥ 0.05, * for P < 0.05, ** for P < 0.01, *** for P < 0.001, and **** for P < 0.0001.

### Inactivation of *Myrf* disrupted mesothelium cell specification

Aside from signaling pathway genes, from the bulk RNAseq, several mesothelial markers (*Upk3b, Gpm6a, Frem2, Wt1)* were among the top 20 differentially expressed genes in the *Tbx4tetOcre;MyrfE6* mutant (Table S1). This raises the possibilities that the changes in gene expression in the *Tbx4tetOcre;MyrfE6* mutant could be the consequence of (1) reduced number of mesothelial cells and/or (2) downregulation of these markers at a per cell basis. To test these two possibilities, we crossed the *Tbx4tetOcre;MyrfE6* mutant to a *R26tdTomat*o reporter (hereafter *Tbx4tetOcre;Myrf;tdTomE6*) to label cre-lineaged cells and perform immunostaining for WT1, as an example of a differentially expressed factor, to assess its protein level per cell (Figure 2C-E). In the cre control, as expected, the majority of lung mesenchymal cells are recombined tdTom+ cells (Fig. 2D)^35^. These include ∼93% of WT1+ mesothelial cells that continuously ensheathe the lung (Figure 2F). In the *Tbx4tetOcre;Myrf;tdTomE6* mutant, however, we observed several phenotypes. First, there are noticeable patches of cells in the mesothelial layer around the lung that are tdTom-, accounting for ∼47% of all WT1+ cells (Figure 2E-F, white arrowheads). Such an increase in tdTom-cells seems to be only in the mesothelial layer, and no increase in mosaicism is observed in the mesenchyme. This observation raised the possibility that non-recombined wild-type cells were recruited due to a deficiency of lung mesothelial cells. Consistent with this, there is a reduced number of recombined (WT1+tdTom+) mesothelial cells per length (um) of the lung perimeter (Figure 2E,G). Even adding on the increased unrecombined cells, the total number of Wt1+ (WT1+tdTom+ and Wt1+tdTom-) mesothelial cells is decreased in the *Tbx4tetOcre;Myrf;tdTomE6* mutant compared to control (Figure 2G). Analysis of cell proliferation and apoptosis by MKi67 and cleaved caspase 3 immunostaining, respectively, at two static stages of E13.5 and E10.5 did not reveal any differences (Figure S2A-E). However, it remains possible that a minor deficiency in either cell proliferation or cell survival could cumulate across stages to lead to reduced mesothelial cell number. Focusing in on tdTom+, recombined mesothelial cells in the *Tbx4tetOcre;Myrf;tdTomE6* lung showed reduced WT1 expression compared to unrecombined (WT1+tdTom-) mesothelial cells in the same embryo (Figure 2E, red arrowheads). These data together suggest that inactivation of *Myrf* compromised mesothelium specification, leading to both reduced mesothelial cell number as well as their expression of mesothelial markers.

### Inactivation of *Myrf* led to increased differentiation into smooth muscle cells

To focus in on *Myrf* requirement in lung mesothelial cells, we inactivated *Myrf* specifically in the mesothelium by mating the *Myrf^Fl/Fl^* conditional strain to the *Wt1^creERT2^* cre driver and *R26R;tdTomato* reporter to generate *Wt1^creERT2^;Myrf^Fl/Fl^;tdTom/+* (hereafter, *Wt1creER;Myrf;tdTomE12*) mutants. We performed inactivation at E12.5, the branching stage of development, and analyzed the mutants at E16.5, when mesothelial cells have progenitor activity and differentiate into multiple mesenchymal cell types^10–14^. Inactivation at E12.5 did not result in a diaphragmatic phenotype (data not shown), allowing us to characterize lung-intrinsic defects in the mutant (Figure 3A-B). *Wt1creER;Myrf;tdTomE12* mutants displayed lung lobes that were smaller and rounded (Figure 3B). Lineage tracing of mesothelial cells revealed that the *Wt1creER;Myrf;tdTomE12* mutant displayed not only a reduction of WT1 expression, but also mesothelial thickening, suggesting an altered differentiation state (Figure 3C).

**Figure 3:**
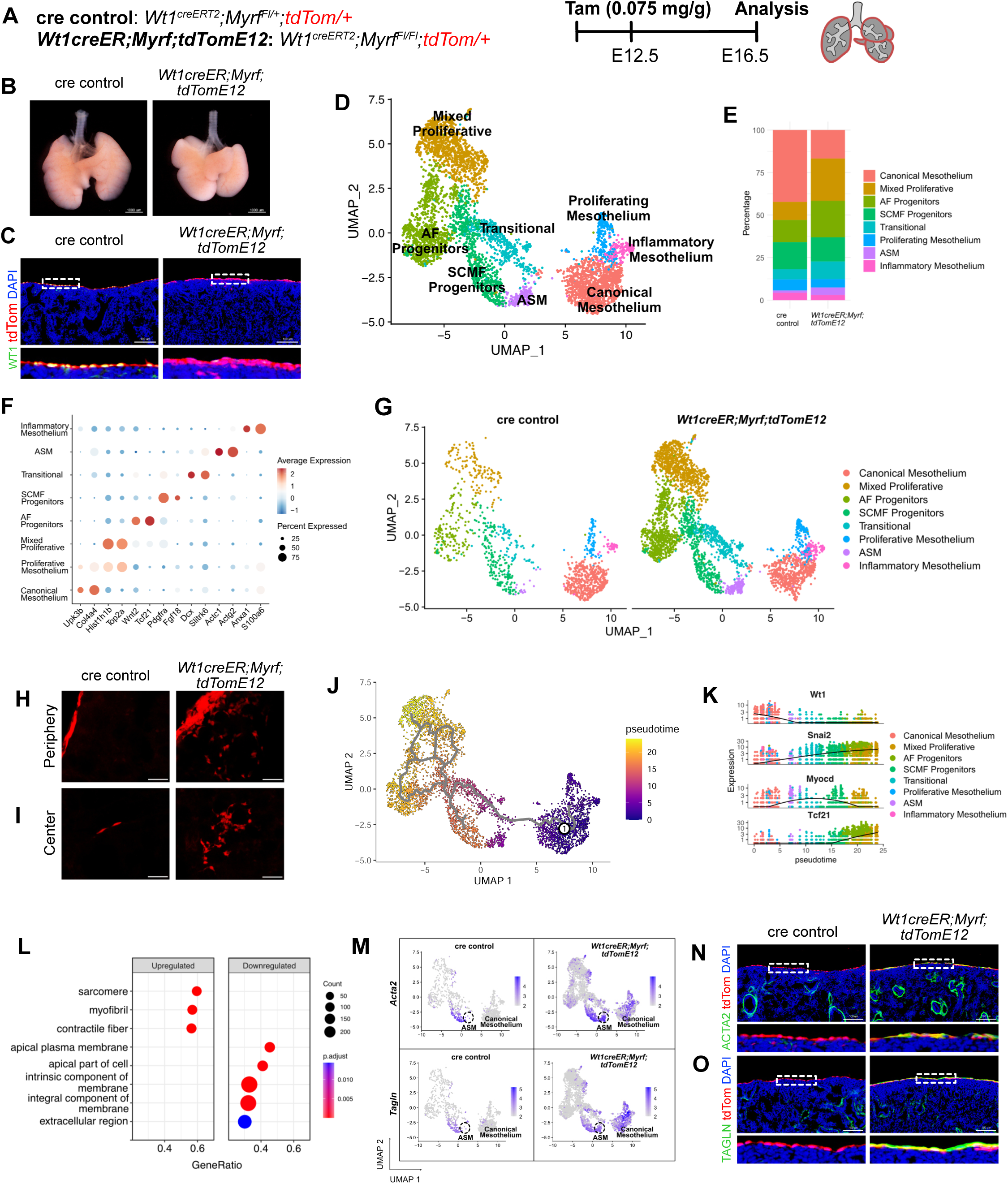
Single-Cell RNA Sequencing of the *Myrf* Mutant Mesothelium. **(A)** Timeline of inactivation and analysis of the *Wt1creER;Myrf;tdTomE12* mutant and cre control. **(B)** Lung whole mount image of *Wt1creER;Myrf;tdTomE12*mutant and cre control. **(C)** Immunostaining for WT1(green) and tdTom(red) in the *Wt1creER;Myrf;tdTomE12* mutant and cre control. **(D)** Integrated UMAP and populations of the *Wt1creER;Myrf;tdTomE12* mutant and cre control at E16.5. **(E)** Proportion of each of the identified populations split by genotype. **(F)** Representative markers of each of the identified populations. **(G)** UMAP split by mutant and control genotype. **(H)** Thick cryosections (99 um) of periphery lung sections of the *Wt1creER;Myrf;tdTomE12* mutant and cre control. **(I)** Thick cryosections (99 um) of center lung sections of the *Wt1creER;Myrf;tdTomE12* mutant and cre control. **(J)** Monocle trajectory analysis of the mesothelium and mesothelium derived populations. **(K)** Expression *of Wt1,Snai2*, *Myocd* and *Tcf21* as a function of pseudotime. **(L)** GSEA enrichment analysis of upregulated and downregulated differentially expressed genes in the *Wt1creER;Myrf;tdTom;E12* mutant. **(M)** Featureplot for *Acta2* and *Tagln.* **(N)** Immunostaining for ACTA2(green) and tdTom(red) in the *Wt1creER;Myrf;tdTomE12* mutant and cre control. **(O)** Immunostaining for TAGLN(green) and tdTom(red) in the *Wt1creER;Myrf;tdTomE12* mutant and cre control.

To further characterize the lineage-traced *Wt1creER;Myrf;tdTomE12* mutant mesothelial cells, we performed single-cell RNA sequencing (scRNAseq) at E16.5 (Figure S3A), pooling sorted tdTom+ cells from multiple embryos for each of the mutant or control group. Among all dissociated lung cells, there was a proportional increase of tdTom+ cells in the mutant compared to control (0.43% vs 1.31%) (Figure S3A-B). After quality control and filtering, a total of 1211 control and 3533 mutant high-quality cells were integrated with Harmony^41^ (Figure 3D- G, Figure S3C-D). Three mesothelium populations were identified both in the mutant and control: Canonical Mesothelium (*Upk3b*+*Col4a4*+), Proliferative Mesothelium (*Hist1h1b*+*Top2a*+) and Inflammatory Mesothelium (*Anxa1*+*S100a6*+). Consistent with reports that mesothelial cells undergo EMT and give rise to lung mesenchymal cell types during development^10–14^, multiple mesenchymal cell types were observed in both the mutant and control *Wt1*-lineage samples. These included *Wnt2*+*Tcf21*+ alveolar fibroblast (AF) Progenitors, *Pdgfra*+*Fgf18*+ secondary crest myofibroblast (SCMF) Progenitors, *Actc1*+*Actg2*+ airway smooth muscle (ASM) cells, *Hist1h1b*+*Top2a*+ Mixed Proliferative cells, and a novel *Dcx*+*Slitrk6*+ Transitional population, that on the UMAP, bridges the mesothelium and some of their derivative populations (Figure 3F-G, TableS2). While the proportion of mesothelial cells decreased in the mutant, the proportion of their derivative mesenchymal cells increased (Figure 3E). Thick cryosections (99 um) of lungs confirmed an increase of migratory tdTom+ cells at both the periphery and center of the mutant lung compared to control (Figure 3H-I).

Next, we used Monocle3 to reconstruct the differentiation trajectory using the mutant and control combined object. Two major branches were identified, one connecting the AF progenitors and Proliferative Mixed population, and one ending at the SCMF Progenitor Population (Fig 3J). Airway smooth muscle cells were not a major end point of the trajectory, likely due to the rarity of mesothelium-lineaged cells that give rise to this population. Indeed, immunostaining for ACTA2 revealed the rare presence of tdTom+ACTA2+ cells along the branching airways (Figure S3E, S3G). The Transitional cells were situated early in the trajectory and at the major branchpoint, consistent with this being a precursor population. Transitional cell marker, DCX, has been shown to be a neuronal microtubule protein important for cell migration^42^. Indeed, mesothelial lineaged cells in the sub-mesothelial region, and to a lesser extent, more centrally in the lung, are DCX+ by immunostaining (Figure S3F, S3H). We next identified genes differentially expressed along the trajectory. As *Wt1* decreased along the trajectory, EMT transcription factor gene *Snai2* increased along the trajectory (Fig 3K, TableS3). Transcription factor genes *Myocd* and *Tcf21* were enriched in the SCMF Progenitor trajectory and AF Progenitor trajectory, respectively. (Fig 3K, TableS3). The finding that mesothelial derived cells show distinct gene expression programs indicate that during development, a subset of mesothelial cells make a clear switch to mesenchymal cell fates.

To address the primary molecular defects that result from loss of *Myrf* in the mesothelium, we performed differential gene expression analysis in the Canonical Mesothelium population in the *Wt1creER;Myrf;tdTomE12* mutant versus control (TableS4). Gene set enrichment analysis (GSEA) of downregulated genes included terms such as apical plasma membrane, intrinsic component of membrane, and extracellular region (Figure 3L). Top downregulated genes included mesothelium-enriched ECM protein genes *Upk3b, Upk1b* and *Gpm6a.* While their function in the lung is unknown, uroplakin proteins are essential components of the urothelial plaques, and in the lung, they may contribute to the mesothelial glyocoalyx^43,44^. Corroborating gene expression changes observed in the *Tbx4tetOcre;MyrfE6* bulk RNAseq dataset, developmental signals and enzymes were downregulated (*Fgf9, Raldh2, Rdh10, Wnt9a, Tgfb3*) (TableS4). These data suggest compromised mesothelial characteristics. GSEA of upregulated genes revealed enrichment of terms such as sarcomere, myofibril, and contractile fiber which is consistent with the smooth muscle genes upregulated in the *Wt1creER;Myrf;tdTomE12* mutant, such as *Acta2, Tagln,* and *Des* (Figure 3L, TableS4). These genes, which are expressed in the smooth muscle cells in the control, displayed robust ectopic expression in the mesothelium of the *Wt1creER;Myrf;tdTomE12* mutant (Figure 3M-O). Dual expression of mesothelial (WT1^low^) and mesenchymal (Acta2^hi^ Tagln^hi^) cell features are indicative of partial EMT. Altogether, these findings suggest that loss of *Myrf* results in increased conversion to mesenchymal cell types starting from the lung periphery.

### Inactivation of *Myrf* disrupted mesothelium polarization

During development, mesothelial cells are known to transition from a cuboidal to a flattened cell morphology^10^. This process is accompanied by the establishment of apical-basal cell polarity, which plays a key role in organizing cell junctions necessary for forming the semi-permeable tissue barrier that protects organs from the external environment^1,8^. From the scRNAseq data, GO category “apical plasma membrane”, which includes polarity proteins, was enriched in downregulated genes in *Wt1creER;Myrf;tdTomE12* mutant mesothelial cells. We next investigated the change of polarity on the protein level by assessing the expression of EZRIN, which is an apical protein that serves as a membrane-cytoskeleton linker^45^. In the *Wt1creER;Myrf;tdTomE12* mutant, we observed reduced EZRIN signal, especially in tdTom+ mesothelial regions that show thickening (Figure 4B). To visualize changes in cell junctions, we performed immunostaining for β-CATENIN, which is known to be associated with adherens junctions at the lateral surface of the epicardium^46^. In the control, β-CATENIN is observed at the lateral regions of the tdTom+ lung mesothelium; however, in the *Wt1creER;Myrf;tdTomE12* mutant, β-CATENIN staining is diffuse (Figure 4C).

**Figure 4:**
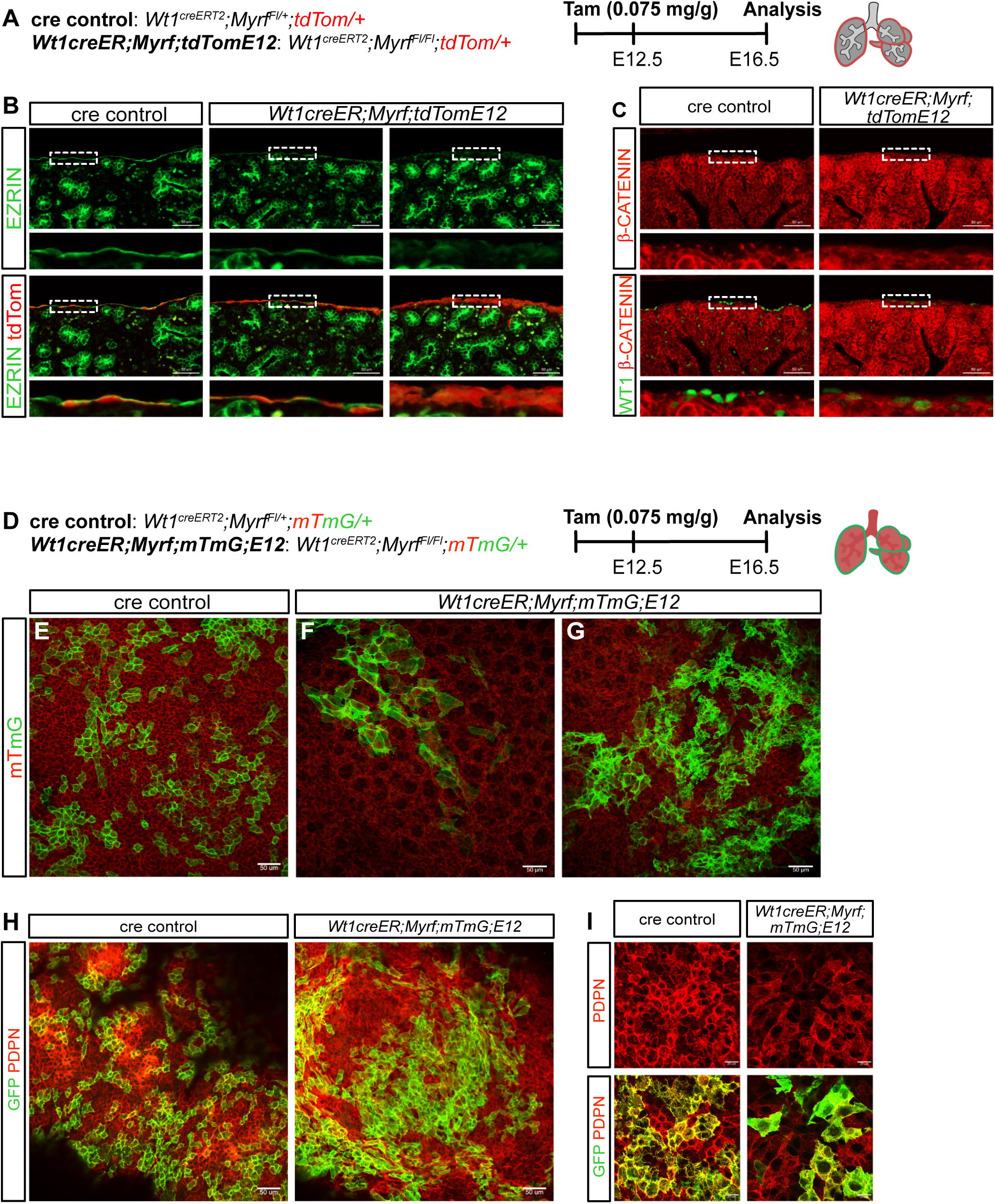
The *Myrf* Mutant Lacks a Single Layered Polarized Mesothelium. **(A)** Timeline of inactivation and analysis of the *Wt1creER;Myrf;tdTomE12* mutant and cre control. **(B)** Immunostaining for EZRIN(green) and tdTom(red) in the *Wt1creER;Myrf;tdTomE12* mutant and cre control. **(C)** Immunostaining for WT1(green) and β-CATENIN(red) in the *Wt1creER;Myrf;tdTomE12* mutant and cre control. **(D)** Timeline of inactivation and analysis of the *Wt1creER;Myrf;mTmG;E12* mutant and cre control. **(E)** Representative whole mount image of the cre control. **(F)** Representative whole mount image of the *Wt1creER;Myrf;mTmG;E12* mutant containing morphology 1. **(G)** Representative whole mount image of the *Wt1creER;Myrf;mTmG;E12* mutant containing morphology 2. **(H)** Immunostaining for GFP(green) and PDPN(red) in the *Wt1creER;Myrf;mTmG;E12* mutant and cre control at 20x. **(I)** Immunostaining for GFP(green) and PDPN(red) in the *Wt1creER;Myrf;mTmG;E12* mutant and cre control at 63x.

To visualize changes in mesothelial cell morphology, we mated the *Myrf^Fl/Fl^* conditional strain to the *Wt1^creERT2^* cre driver and *R26;mTmG* reporter to generate *Wt1^CreERT2^;Myrf^Fl/Fl^;mTmG/+* mutants (hereafter, *Wt1creER;Myrf;mtmG;E12*, Figure 4D). This reporter line has membrane-localized tdTomato expression that switches to membrane-localized GFP expression upon cre recombination^47^. In the control, with the mTmG reporter and 0.075 mg/g dose of tamoxifen administered at E12.5 and assayed at E16.5, the *Wt1^creERT2^* cre driver was able to recombine the reporter in ∼30-40% of the mesothelial cells. We observed groups of squamous GFP+ mesothelial cells connect into a monolayer on the surface of the lung (Figure 4E). In contrast, in the mutant, GFP+ cells show two types of morphologies. In regions that remained monolayer, GFP+ cells in the mutant were larger than in the control, as if they were stretched to cover the surface of the lung (Figure 4F). In regions that are thickened into multiple cell layers, GFP+ cells were highly disorganized and adopted a spindle-shaped, elongated morphology, consistent with fibroblast/smooth muscle cell characteristics (Figure 4G). Immunostaining for PDPN, which at this stage (E16.5) labels cells with epithelial characteristics, including mesothelial cells on the lung surface, confirmed that in the mutant, these GFP+ cells retain some mesothelial/epithelial characteristics (Figure 4H-I). These imaging results align with the single cell transcriptomic data and together suggest that mutant mesothelial cells have acquired a hybrid differentiation state.

### Inactivation of *Myrf* after birth led to mesothelial thickening without ectopic differentiation into ACTA2+ cells

To determine the role of *Myrf* after birth and post establishment of the mesothelium, we utilized the *Wt1creER;Myrf;tdTom* mutant and performed inactivation of *Myrf* using tamoxifen treatment starting at P1 (hereafter, *Wt1creER;Myrf;tdTomP1*) (Figure S4A). The gross lung size and shape of the cre control and *Wt1creER;Myrf;tdTom* mutant lung were similar at P39, however, mutants exhibited apparent mesothelial thickening (Figure S4B, arrow). Examination by sectioning revealed mesothelial thickening in the postnatal *Wt1creER;Myrf;tdTomP1* mutant, similar to that in the prenatal mutant. However, immunostaining for ACTA2+ showed that unlike the prenatal mutant, cells in the thickened region of the postnatal mutant do not express ACTA2, suggesting that that these cells did not convert into smooth muscle (Figure S4C). Interestingly, also unlike the prenatal mutant, immunostaining for WT1 revealed an increase rather than a decrease in WT1 expression compared to control, suggesting that these cells have maintained a mesothelial characteristic (Figure S4D).

### H3K27ac CUT&RUN revealed MYRF as a regulator of mesothelium cell fate

To systematically address the transcriptional control of the developing mesothelium, we performed a CUT&RUN assay for activated enhancer-associated H3K27ac histone in wild-type E16.5 lung mesothelium and epithelium from the same lungs for comparison (n=2 biological replicates) (Figure 5A-B). To enrich for lung mesothelium versus epithelium, we developed a flow cytometry panel, using EPCAM-PDPN+CD45-CD31- surface staining to enrich for the mesothelium and EPCAM+PDPN+CD45-CD31- surface staining to enrich for the epithelium. This staining panel was validated using E16.5 *Wt1creER;Myrf;tdTomE12* control lungs where >80% of EPCAM-PDPN+CD45-CD31- cells were tdTom+, suggesting substantial enrichment of the mesothelium population (Figure S5A-B).

**Figure 5:**
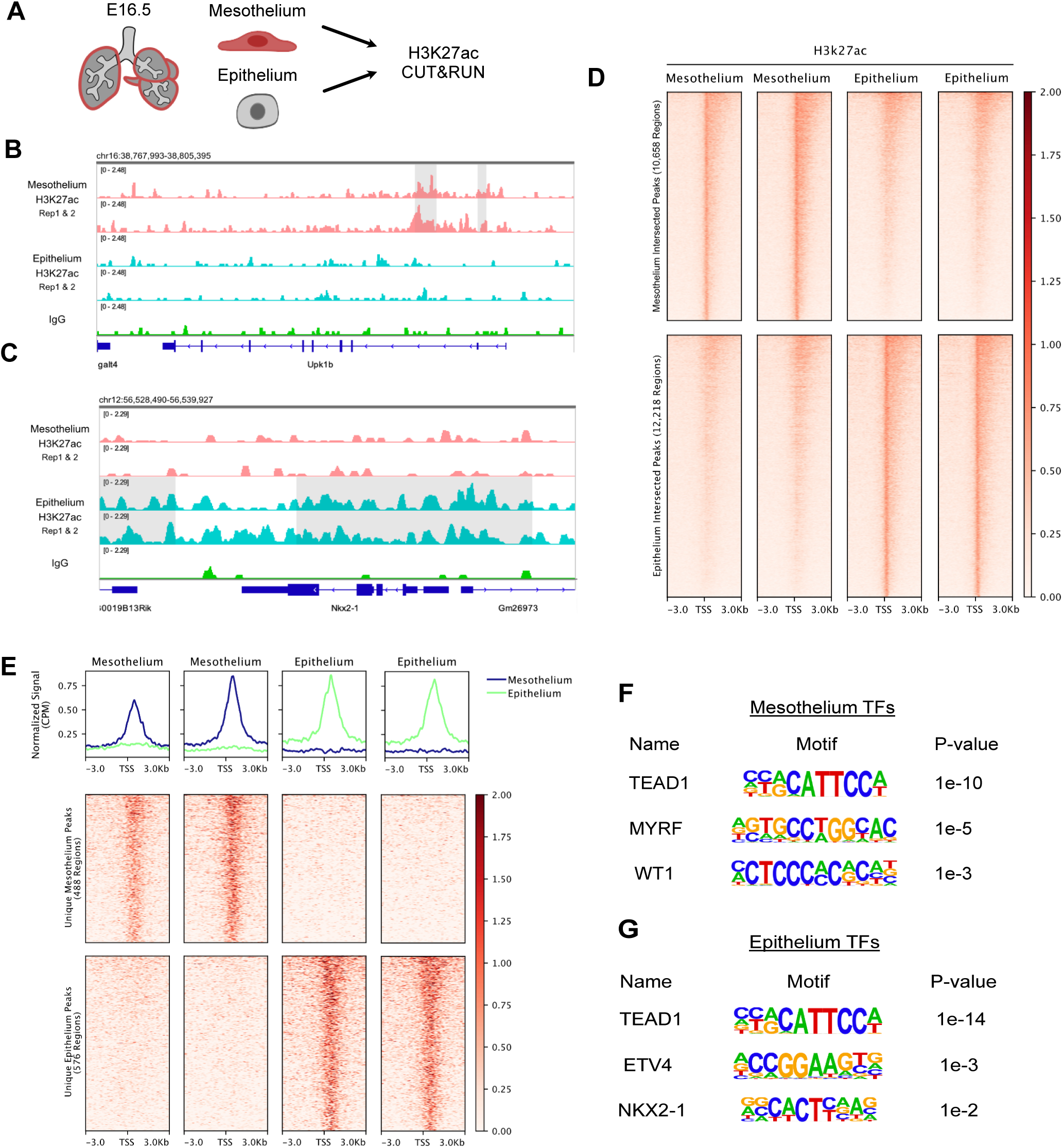
E16.5 Mesothelium and Epithelium H3K27ac CUT&RUN. **(A)** E16.5 mesothelium and epithelium CUT&RUN experiment diagram. **(B)** Mesothelium and Epithelium H3K27ac and IgG peaks at the Upk1b gene locus. **(C)** Mesothelium and Epithelium H3K27ac and IgG peaks at the NKx2-1 gene locus. **(D)** H3K27ac peaks in the mesothelium and epithelium across intersected peak sets centered at the TSS and flanked by +/- 3 kb. **(E)** Differentially enriched H3K27ac peaks in the mesothelium and epithelium centered at the TSS and flanked by +/- 3 kb. **(F)** Homer motifs identified in differentially enriched peaks in the mesothelium. **(G)** Homer motifs identified in differentially enriched peaks in the epithelium.

A total of 10,658 and 12,218 reproducible peaks were identified in mesothelial and epithelial cells, respectively (Figure 5B-D). This included regions near genes *Upk1b*, enriched in the mesothelium, and Nkx2-1, enriched in the epithelium. Annotation of the peaks demonstrated that 39.9% and 29.7% of peaks reside within +/- 1kb of promoters in the mesothelium dataset and the epithelium dataset, respectively, suggesting that most peaks are in the distal regions (Figure S5C). Comparing the mesothelium to epithelium, 1,064 peaks were differentially enriched (TableS5, Figure 5E, S5D). MYRF emerged from motif analysis as a top transcription factor predicted to bind to mesothelium enriched peaks, together with TEAD1 and WT1 (Figure 5F). In comparison, TEAD1, ETV4, and Nkx2-1 emerged as top transcription factors predicted to bind to epithelium enriched peaks (Figure 5G). We note that TEAD1 motifs were enriched in both cell types, consistent with the diverse roles of Hippo-Yap signaling in multiple cell types in lung development and repair^48^.

Next, we interrogated MYRF binding sites across the 488 differentially enriched peaks in the lung mesothelium. 27% of peaks had MYRF motifs and 7.8% of peaks had a MYRF motif associated with a gene that was differentially expressed in the *Wt1creER;Myrf;tdTomE12* mutant mesothelium (Figure 3). These MYRF-containing mesothelial peaks included regions within downregulated genes, such as *Upk1b*, *Cdh11*, and *Nod1*, as well as upregulated genes, such as *Meis1* and *Hoxb5* (TableS5). These results further support the transcriptional role of MYRF in the lung mesothelium.

### *Yap*/*Taz* function in parallel with *Myrf* to promote lung mesothelium cell fate

Given that TEAD1 motifs are enriched in the mesothelial peaks, we next addressed if YAP/TEAD function in these cells and how they may intersect with MYRF. In the *Wt1creER;Myrf;tdTomE12* mutant and control, nuclear YAP immunostaining signal was observed throughout the lung, including the lung mesothelium (Figure 6A-B). To investigate the role of Hippo-YAP/TAZ signaling in the lung mesothelium, we generated mesothelial *Taz* (*Wt1^creERT2^;Myrf^Fl/+^;Yap^Fl/+^;Taz^Fl/Fl^*, hereafter *Taz* mutant) and *Yap/Taz (Wt1^creERT2^;Myrf^Fl/+^;Yap^Fl/Fl^;Taz^Fl/Fl^,* hereafter *Yap;Taz* mutant) mutants (Figure 6C). To address if mutant mesothelial cells have converted into smooth muscle cells, we performed immunostaining for smooth muscle marker ACTA2 and quantified positive pixels at the lung periphery. *Yap;Taz* mutants, but not *Taz* mutants, displayed a trending increase in ACTA2+ pixels at the lung periphery, suggesting a similarity in this phenotype to *Myrf* mutants (Figure 6F-G, J).

**Figure 6:**
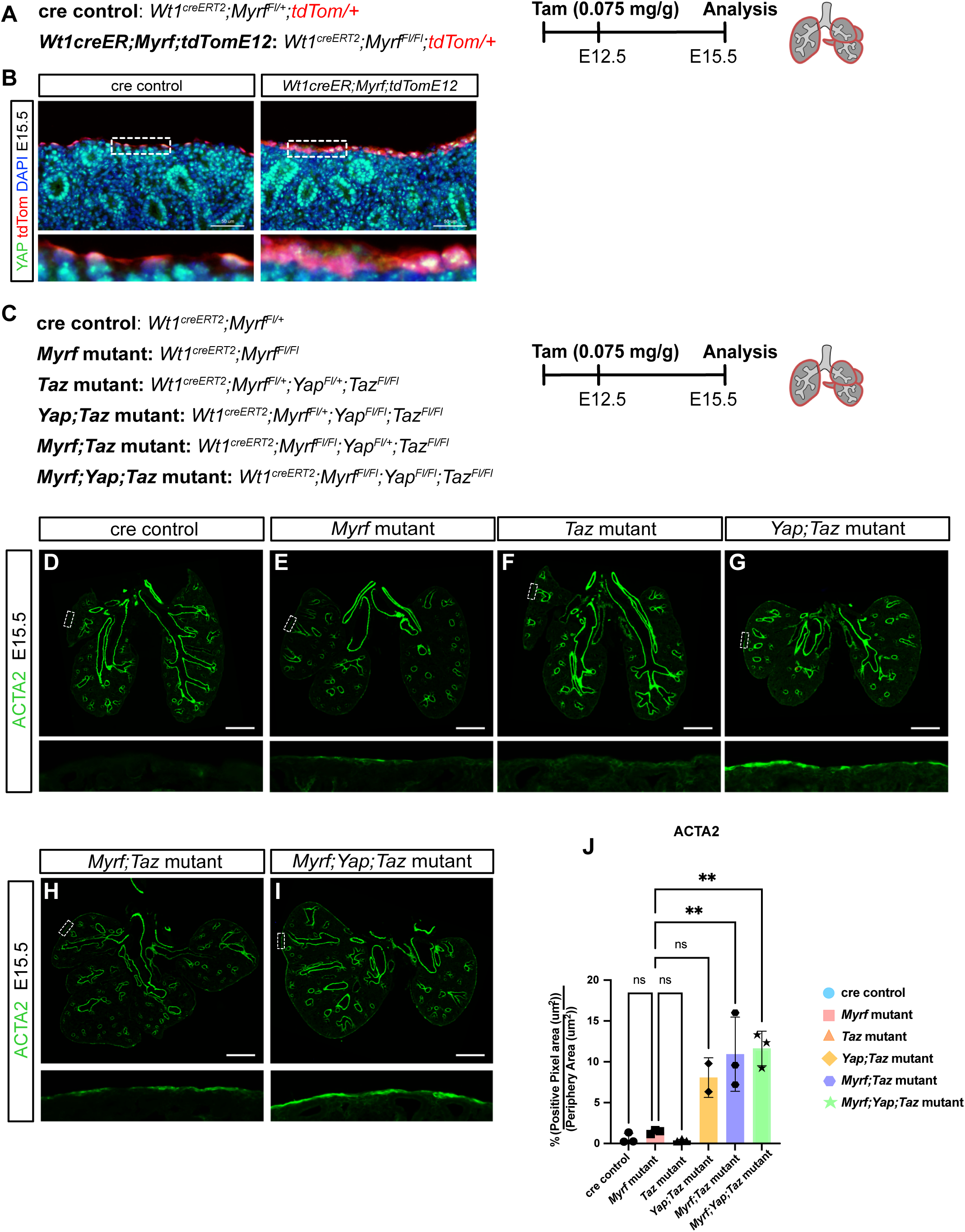
Hippo-Yap/Taz Signaling in the Lung Mesothelium. **(A)** Timeline of inactivation and analysis of the *Wt1CreER;Myrf;tdTomE12* mutant and cre control. **(B)** Immunostaining for YAP(green) and tdTom(red) in the E15.5 *Wt1CreER;Myrf;tdTomE12* mutant and cre control. **(C)** Timeline of inactivation and analysis of the cre control, *Myrf*, *Taz, Yap;Taz, Myrf;Taz*, and *Myrf;Yap;Taz* mutants. **(D)** Representative ACTA2(green) immunostaining of the cre control. **(E)** Representative ACTA2(green) immunostaining of the *Myrf* mutant. **(F)** Representative ACTA2(green) immunostaining of the *Taz* mutant. **(G)** Representative ACTA2(green) immunostaining of the *Yap;Taz* mutant. **(H)** Representative ACTA2(green) immunostaining of the *Myrf;Taz* mutant. **(I)** Representative ACTA2(green) immunostaining of the *Myrf;Yap;Taz* mutant. **(J)** Quantification of ACTA2 positive pixel area over the periphery area in the cre control, *Myrf*, *Taz, Yap;Taz, Myrf;Taz*, and *Myrf;Yap;Taz* mutants. *Myrf* mutant interactions are displayed in the graph. One-way ANOVA Tukey’s multiple comparison test was used for 6J. ns (not significant) for P ≥ 0.05, * for P < 0.05, ** for P < 0.01, *** for P < 0.001, **** for P < 0.0001.

To determine the genetic interaction between *Myrf* and *Yap/Taz* in the lung mesothelium, we inactivated *Taz* (*Wt1^creERT2^;Myrf^Fl/Fl^;Yap^Fl/+^;Taz^Fl/Fl^*, hereafter *Myrf;Taz* mutant) and both *Yap* and *Taz* (*Wt1^creERT2^;Myrf^Fl/Fl^;Yap^Fl/Fl^;Taz^Fl/Fl^*, hereafter *Myrf;Yap;Taz* mutant ) in the *Myrf* mutant background (Figure 6C,H-I). A statistically significant increase of ACTA2+ pixels was observed in the *Myrf;Taz* mutant and *Myrf;Yap;Taz* mutant, with each compared to the *Myrf* single mutant (Figure 6D-J). Immunostaining for smooth muscle TAGLN was also performed across all mutants, and while a trending increase in TAGLN immunostaining was observed in the *Yap;Taz*, *Myrf;Taz,* and *Myrf;Yap;Taz* mutants, no comparison was statistically significant due to greater variation compared to ACTA2 staining (Figure S6A-G). To quantify more precisely, we incorporated a tdTom reporter in the *Myrf;Taz* and *Myrf;Yap;Taz* mutants (Figure S6H). Quantification of TAGLN+ and ACTA2+ pixels in the tdTom+ regions revealed a statistically significant increase (Figure S6I-N). Altogether, these data are consistent with the possibility that MYRF and YAP/TAZ function in parallel to control mesothelial cell differentiation.

### Ectopic peripheral smooth muscle cells persist but do not progress in the adult lung of the *Myrf* mutant

In IPF lungs, ACTA2+ cell accumulate in the subpleural fibroblasts, and have been postulated to play a role in the stereotypic pleural-to-center pattern of fibrosis progression^49,50^. We next addressed if the ectopic ACTA2+ cell phenotype in the *Myrf* development mutants persists and/or progresses in the postnatal lung. Since *Wt1creER;Myrf;tdTomE12* mutants die at birth (data not shown), possibly due to inactivation of *Myrf* in other tissues (e.g. heart and liver), we returned to the *Tbx4tetOcre;Myrf* (*Tbx4-rtTA/+;tetO-cre/+;Myrf^Fl/Fl^)* mutant as such inactivation is primarily in the lung mesenchyme and diaphragm. To bypass herniation, which caused lethality of *Tbx4tetOcre;MyrfE6* mice, we inactivated *Myrf* at E12.5 (hereafter, *Tbx4tetOcre;MyrfE12*) (Figure 7A, S7A). No herniation was observed, and the mice survived postnatally. At P22 the *Tbx4tetOcre;MyrfE12* lung was misshapen and the separation of lobes, in particular the cranial and caudal lobes, were frequently incomplete (Figure 7B, S7A). H&E staining showed increased thickening of the mesothelium comparing P22 to P1 (Figure 7C). Immunostaining for club (SCGB1A1), ciliated (FOXJ1), AT1 (HOPX), and AT2 (SFTPC) cell markers revealed normal differentiation of airway and alveolar epithelial cells, suggesting normal alveolar and airway cell architecture (Figure S7B-C).

**Figure 7:**
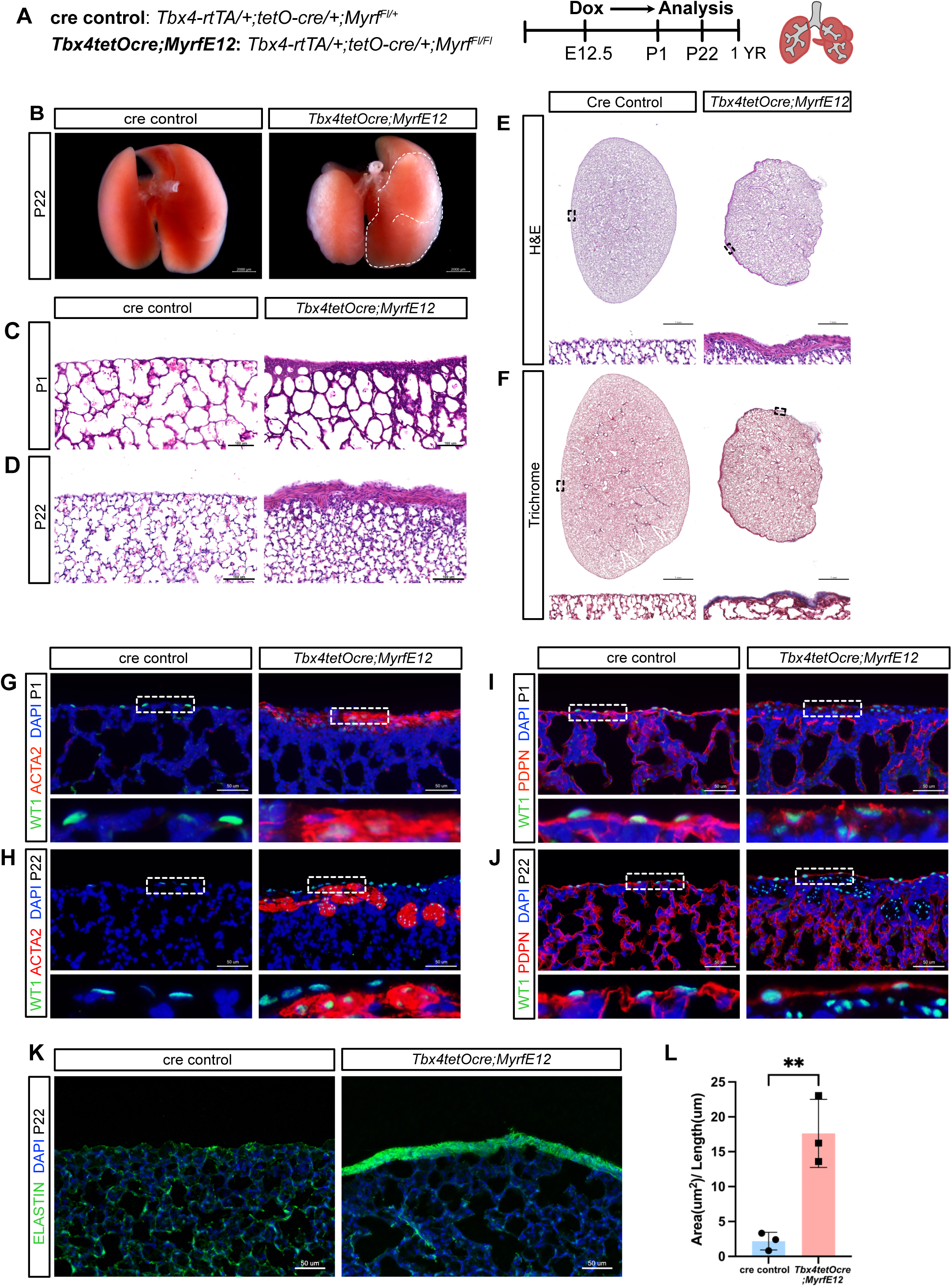
Postnatal Analysis of the *MyrfE12Cko* Mutant Lung. **(A)** Timeline of inactivation and analysis of the *Tbx4tetOcre;MyrfE12* mutants and cre controls. **(B)** Representative whole mount images of *Tbx4tetOcre;MyrfE12* mutants and cre controls at P22. **(C)** H&E images of *Tbx4tetOcre;MyrfE12* mutant and cre control at P1. **(D)** H&E images of *Tbx4tetOcre;MyrfE12* mutant and cre control at P22. **(E)** Slide scanned H&E image of the *Tbx4tetOcre;MyrfE12* mutant and cre control at P22. **(F)** Slide scanned Trichrome image of the *Tbx4tetOcre;MyrfE12* mutant and cre control at P22. Blue=collagen. **(G)** Immunostaining for WT1(green) and ACTA2(red) in the *Tbx4tetOcre;MyrfE12* mutant and cre control at P1. **(H)** Immunostaining for WT1(green) and ACTA2(red) in the *Tbx4tetOcre;MyrfE12* mutant and cre control at P22. **(I)** Immunostaining for WT1(green) and PDPN(red) in the *Tbx4tetOcre;MyrfE12* mutant and cre control at P1. **(J)** Immunostaining for WT1(green) and PDPN(red) in the *Tbx4tetOcre;MyrfE12* mutant and cre control at P22. **(K)** Immunostaining for ELASTIN(green) in the *Tbx4tetOcre;MyrfE12* mutant and cre control at P22. **(L)** Quantification of ELASTIN thickness at the lung periphery (Area(um^2^)/Length(um)).

Similar to the *Wt1CreERT2;Myrf* mutant, an increase in ACTA2+ cells were detected at the mesothelium and sub-mesothelial regions of the *Tbx4tetOcre;MyrfE12* lung at P1 (Figure 7G). By P22, ACTA2+ cells formed smooth muscle bundles at the lung periphery and were devoid of PDPN (Figure 7H-J). Immunostaining for WT1 revealed a minor reduction at P1, consistent with observation at prenatal stages. But by P22, WT1 expression has returned both in the mesothelium and some of the ACTA2+ cells in the sub-mesothelial regions (Figure 7G-H). To verify that the WT1 expression occurred in cells that have undergone recombination, we incorporated cre activity reporter *R26;TdTomato* into the *Tbx4tetOcre;MyrfE12* mutant background (hereafter *Tbx4tetOcre;Myrf;tdTomE12*). There were many TdTom+ cells that expressed Wt1+ in the thickened sub-mesothelial region in the *Tbx4tetOcre;Myrf;tdTomE12* mutant (Figure S7D). These data together suggest that by weaning, WT1 expression is no longer dependent on *Myrf*, similar to the *Wt1creER;Myrf;tdTomP1* mutant (Figure S4D).

We traced the *MyrfckoE12* mutant postnatal phenotype to 1 year. The thickening ACTA2+ lesions persisted but stayed confined to the periphery of the lung in the *Tbx4tetOcre;MyrfE12* mutant, unlike the progressive nature of myofibroblasts from periphery to center in IPF (Figure S7E). Trichrome staining at P22 showed minor, but consistent collagen accumulation at the periphery of the *Tbx4tetOcre;MyrfE12* mutant lung (Figure S7F). ELASTIN staining at P22 showed a significant accumulation at the periphery of the *Tbx4tetOcre;MyrfE12* mutant lung (Figure 7K-L). Such presence of non-progressive collagen and elastin accumulation are features of pleuroparenchymal fibroelastosis (PPFE), a rare lung fibrotic disease with no known cause.

## Discussion

Across all organs, the mechanisms underlying mesothelium development and homeostasis are poorly understood. In this study, using the lung mesothelium as a setting, we identified MYRF as a key transcriptional regulator of mesothelial cell fate. Inactivation of *Myrf* at the start of lung development impaired mesothelial cell specification and compromised their role as a signaling center for lung growth. Inactivation at the branching stage impaired mesothelial cell polarity, resulting in partial EMT and the adoption of smooth muscle cell characteristics. This unique, thickened, elastin-expressing ectopic smooth muscle layer persists into the adult, leading to a phenotype that, at the histological level, shows a resemblance to PPFE. While the cause of PPFE remains unclear and it is not known whether disruption of *MYRF* is involved, our findings establish the *Myrf* mutant as a potential mouse model to gain insights into this disease. In addition to *Myrf,* our data demonstrated that *Yap/Taz* are also critical regulators of mesothelium cell differentiation. Together, the MYRF, WT1 and YAP/TAZ transcription factors establish a core transcription factor regulatory network that controls mesothelial cell development.

With the mesothelium encasing internal organs, it makes sense that it serves as a cardinal regulator of organ growth. In support of this role, we show that in the *Tbx4tetOcre;MyrfE6* mutant, defective mesothelial cell specification led to smaller lungs, which is apparent prior to their compression by herniated abdominal organs. It is worth noting that in human patients that carry variants in *MYRF*, pulmonary hypoplasia has also been observed in a subset of patients without diaphragmatic abnormalities and herniation^27,31^. Data from both bulk, as well as single cell RNAseq, showed that regardless of the timing of *Myrf* inactivation, either at the start of lung development, or at branching stage, mutants showed reduced expression of genes in the FGF, retinoic acid, and other growth factor pathways. Concomitantly, there is a decrease in mesothelial cell number in the *Tbx4tetOcre;MyrfE6* mutant. Reduced growth factor expression, due to both reduced mesothelial cell number and reduced expression per cell, could be the cause for lung hypoplasia. In addition, whether reduced mesothelial cells alter biomechanical constraints of the lung remains to be investigated. Together, these results establish MYRF as a key factor that controls mesothelial cell specification.

Our scRNAseq data of *Wt1creER*-lineaged cells provide the first systematic molecular-level demonstration that in the normal developing lung, mesothelial cells can give rise to alveolar fibroblasts, SCMFs, and smooth muscle cells. These results establish mesothelial cells as a unique multipotent progenitor during development. We further show that MYRF plays a crucial role in the maintenance and execution of such progenitor property. In the absence of *Myrf*, mesothelial cells show increased differentiation into progeny cell types at the expense of self-renewal. In the mutant, even cells that remained at the lung mesothelium have taken on characteristics of smooth muscle. Non-lineaged cells near the mesothelium are also found to ectopically express ACTA2, suggesting a non-cell autonomous effect through either altered signaling or mechanical stress. These results establish MYRF as a key factor that controls mesothelial cell differentiation.

As the outermost cell layer of the lung, the mesothelium is subject to mechanical tension with cyclic breathing movements. Although AT1 cells in the lung and mesothelial cells share a squamous cell morphology, they are from different lineages — AT1 cells are from the endoderm and mesothelial cells are from the mesoderm. In our study, H3K27ac mapping of the mesothelium showed an enrichment of both TEAD and MYRF motifs, while only TEAD, but not MYRF motif was enriched in the epithelium. The selected enrichment of MYRF is corroborated by our functional evidence that *Myrf* is required in the mesothelium, but not in the epithelium, including AT1 cells. We also found that inactivation of *Yap/Taz* on its own led to ectopic ACTA2 expression at the mesothelium. Furthermore, the phenotype is enhanced in the *Myrf* mutant background, suggesting a synergistic relationship in controlling mesothelial cell differentiation. Previous work has shown that mechanical tension promotes AT1 cell fate through regulating nuclear translocation of YAP^51,52,53^. Thus, it is conceivable that like in AT1 cells, mechanical tension may also have a role in modulating mesothelial cell fate.

The persistence of the ectopic smooth muscle cells on the mesothelium surface in the postnatal *Myrf* mutant lung established a striking phenotype of the lung encased in a smooth muscle shell. Ectopic smooth muscle has also been observed in the lung of mesothelial *Ezh2* mutants^54^. Our study provides evidence that that loss of lineage-determining transcription factors can lead to aberrant conversion to other cell fates. We note that based on scRNAseq data, these ectopic *Acta2*+ cells do not express fibrotic signals or markers such as *Cthrc1*, a top marker for fibrotic myofibroblasts^55^. These ACTA2+ cells also do not progress from periphery to center, possibly because they are not fibrotic myofibroblasts. In addition, there is also no senescence associated protein (SASP) signals from damaged alveolar cells that have been shown to be required for fibrosis progression^56^.

While our study focuses on *Myrf* function in lung, given the broad expression of *Myrf* in the mesothelium of other organs and body cavities, our findings lay the foundation for understanding the wide-ranging roles of *Myrf* in organogenesis. In support of this, cardiac and urogenital defects are also common in patients with *MYRF* deficiency^31^. The *Myrf* mutant phenotypes demonstrate it also a causal factor for CDH. Our findings therefore shine the spotlight on the often-overlooked mesothelium, and establishes it as a central player in development, homeostasis and disease pathogenesis.

## Materials and Methods

### Experimental Model and Subject Details

*Myrf^Fl/Fl^, Myrf Δ, Yap^Fl/Fl^Taz^Fl/Fl^,Tbx4-rtTA/+*, *tetO-cre/+, Wt1^CreERT2^, Shhcre^+/-^, Prrx1-cre, R26- tdTomato* and *R26-mTmG* alleles have been described previously^32,47,57,35,58,59,60,61,62^. Tamoxifen was diluted 10:1 corn oil: ethanol and administered intraperitoneally (0.075 mg/g Body weight) at E12.5. Doxycycline was diluted in 1XPBS and administered intraperitoneally (2 mg) at E6.5 or E12.5. Pregnant females injected with Doxycycline were placed on a continuous Doxycycline Diet (625 mg/kg) until takedown. All procedures were performed in accordance with the Institutional Animal Care and Use Committee (IACUC) at the University of California San Diego.

## Method Details

### Tissue Preparation and Immunostaining

Embryos (E13.5), embryonic lungs (E12.5 – E16.5) and postnatal lungs were fixed in 4% Paraformaldehyde (Electron Microscopy Sciences) overnight at 4C degrees. Samples were subsequently processed for paraffin embedding and sectioned at 5 μm. For cryosection, samples were immersed in 30% sucrose for 48 hours, embedded in OCT(Sakura) and sectioned at 10 - 99 μm.

H&E and Trichrome (Abcam) staining was performed using standard procedures. MLI quantification was performed using the Measure MLI Plugin for ImageJ v1.52^63^. A minimum of three evenly spaced lung sections was used from each biological replicate.

Immunostaining was performed using standard procedures. For YAP and pYAP(Ser 127) immunostaining, the Tyramide 488 superboost Kit (Invitrogen) was used. Dapi (1:1000, Sigma) was stained for 5 minutes at room temperature. Following immunostaining, autofluorescence was quenched using Trueblack (Biotium). Slides were mounted using Fluoromount-G mounting media (Invitrogen) and photographed using the Zeiss AxioImager.A2 microscope.

### Whole Mount Immunostaining and Imaging

E13.5 and E16.5 Embryonic lungs were fixed in 4% Paraformaldehyde (Electron Microscopy Sciences) overnight at 4C degrees. E13.5 lungs were dehydrated with a methanol/PBS + 0.1%Tween series (25%, 50%, 75%, 100%) and stored in the −20C. Prior to immunostaining, samples were rehydrated to PBS + 0.1%Tween. Samples were blocked with 10% Goat Serum for two hours at room temperature and incubated overnight with aSMA-Cy3 (Sigma, 1:250) and rabbit anti E-CADHERIN (CST, 1:200) primary antibodies at 4C degrees. The following day, samples were stained with a goat anti-rabbit secondary overnight (Jackson ImmunoResearch; 1:250). Samples were mounted using citifluor (Electron microscopy sciences) and photographed using the Leica SP8 Confocal microscope.

For E16.5 lungs, post-fixed samples were cleared with CUBIC clearing reagent R1 for 3- 5 days. For visualizing endogenous reporter fluorescence, samples were washed with PBS for 6 hours and mounted. For immunostaining, samples were blocked with 10% Goat Serum for two hours at room temperature and incubated with a chicken anti-GFP antibody (Abcam, 1:200) and Syrian hamster anti-PDPN (Developmental Studies,1:100) for three days at room temperature. Samples were stained with goat anti-Chicken 488 (Jackson ImmunoResearch, 1:200) and goat anti-Syrian Hamster Cy3 (Jackson ImmunoResearch, 1:200) for three days at room temperature. Samples were mounted with citifluor (Electron microscopy Sciences) and photographed using the Leica SP8 Confocal microscope.

### RNAscope

Embryos (E13.5) were fixed in 10% NBF for 24 hours at room temperature and processed for paraffin embedding. *Myrf* was detected using RNAscope Probe-Mm-Myrf-C1 (ACDBio) and the RNAscope Multiplex Fluorescent V2 assay (ACDBio). Staining was performed using manufacturers protocol for FFPE.

### Bulk RNA-Seq

E13.5 *Tbx4tetOcre;MyrfE6* mutant and cre control lungs were placed in TRIzol LS and stored in the −80C. Samples were homogenized using a tissuelyser (Qiagen) and RNA was isolated using standard procedures. RNA was purified using the RNAeasy micro Kit (Qiagen). cDNA Libraries were constructed using the Illumina mRNA Stranded Prep (Illumina) and sequenced on the Illumina NovaSeq 6000 instrument, with 50 bp paired end reads. FASTQ files were aligned to the GRCm38 mouse reference genome using STAR^64^. FeatureCounts was used for read counting and DESeq2 was used to perform differential expression^65,66^. Z-Scores of select differentially expressed genes were visualized with R package pheatmap (1.0.12). N=4 samples were sequenced for each genotype and each N represents two pooled samples of the same genotype.

### qRT-PCR

RNA was isolated as described above. cDNA was generated using the iScript cDNA Synthesis Kit (Bio-rad). qRT-PCR was performed using the iTaq Universal SYBR Green Supermix (Bio-rad) and qRT-PCR reactions were run on the Bio-Rad CFX connect real-time PCR machine (Bio-rad). QPCR primers used are listed in TableS6. At least 3 technical replicates and 3 biological replicates were performed per gene target. Cq values were normalized to the Beta-Actin housekeeping gene and statistical significance was performed using Graphpad Prism (10.2.3).

### Tissue Dissociation and Flow Cytometry

For Single Cell RNA-Sequencing, timed *Myrf Fl/Fl;R26TdTomato/R26TdTomato* x *Wt1creERT2;Myrf Fl/Fl* mating’s were set up and pregnant females were injected with Tamoxifen I.P at E12.5. Females were euthanized at E16.5 and mutant and Cre+ embryos were screened for TdTom+ fluorescence. Mutant and cre control lungs were dissected and genotyped based on visual lung/cell shape phenotypes. All genotypes were later confirmed with PCR. Mutant and cre control lungs were pooled (3-5 per genotype) and lung lobes were cut with scissors. Lungs were enzymatically digested for 30 mins at 37C degrees in 5 mL of dissociation media containing RMPI+ (10mM HEPES, 1mM MgCl2, 1mM Cacl2) + 5% FBS, 5U/mL Dispase (Corning), 0.5 mg/mL Collagenase D (Sigma), and 0.05 mg/mL Dnase I (Sigma). Samples were neutralized with RPMI + 10% FBS and filtered with a 40uM strainer (Macs). RBC lysis was performed with 1 mL of ACK lysis buffer (Thermo) for 1 minute at room temperature. Samples were neutralized with DPBS+ 0.2% FBS and filtered with a 40uM strainer. All cell centrifugations were performed at 1500 RPM 4C degrees for 5 minutes. Prior to sorting, samples were resuspended in DPBS+ 0.2% FBS and DAPI (1:16,000). Samples were sorted using the BD FACSAria II (BD Biosciences) and gating was set up using single color controls. Approximately 16,000 cells for each genotype were loaded into the 10x Chromium Controller (10x Genomics).

For CUT&RUN, timed *Myrf Fl/Fl* mating’s were set up. E16.5 females were euthanized, and embryonic lungs were dissociated as described above. After dissociation, cells were resuspended at a max staining volume of 50 million cells/mL. Cells were blocked with BD FC block (1:200, BD Biosciences) for 20 minutes on ice. Cells were stained with the following antibodies: APC/Cyanine7 anti-mouse CD326 (1:500, Biolegend), APC anti-mouse Podoplanin (1:500, Biolegend), FITC anti-mouse CD31 (1:50, Biolegend), Brilliant Violet 510™ anti-mouse CD45 (1:100) for 25 minutes on ice. Two washes with DPBS+ 0.2% FBS were performed, and prior to sorting, samples were resuspended in DPBS + 0.2% FBS and DAPI(1:16,000). Samples were sorted using the BD FACSAria II (BD Biosciences). Mesothelial cells were sorted as PDPN+EPCAM-CD45-CD31-DAPI- and Epithelial cells were sorted as PDPN+EPCAM+CD45- CD31-DAPI-. Approximately 60-100k Mesothelial cells were collected and approximately 100- 500k Epithelial cells were collected.

### Single Cell RNA-Sequencing and Analysis

Single cell RNA Sequencing was performed using the Chromium Single Cell 3’ V3 Kit (10x Genomics). Libraries generated were sequenced on the NovaSeq 6000 at UC San Diego IGM at the school of medicine. FASTQ files were processed with Cell Ranger 6.0.2 (10x Genomics) and reads were aligned with the mm10-2020-A transcriptome to generate the filtered feature barcode matrix.

Mutant and control single cell runs were merged in respect to genotype and processed with R package Seurat (Version 4.3.0.1)^67^. Low quality and duplicate cells were removed with the following parameters: nFeature >2500 & nFeature < 7500, percent.mt <5. Clusters were visualized with Seurat command Dimplot() UMAP projection. Prior to integration low quality populations and populations with mixed identities were removed (populations containing more than one of the following markers: *Sftpc, Hbb-bs, Hopx, Epcam, Nkx2-1*).

Mutant and control samples were integrated with Harmony (Version 1.2.0). To reduce cell-cycle effects, a cell cycle difference regression was performed. A cell-cycle scoring was applied using the cell cycle markers loaded with Seurat (from Tirosh et al, 2015) and the difference between G2M and S phase scores was regressed. Small populations (Approximately 100 cells) were removed and desired populations were subset and re-clustered. Population markers were identified using Seurat FindAllMarkers() command and differential expression testing was performed using Seurat FindMarkers() command. Seurat visualization commands (Dotplot(), Featureplot(), DoHeatmap()) were used to visualize gene expression differences.

Trajectory analysis was performed using Monocle3 (Version 1.2.9) and the SeuratWrappers(version 0.3.1) R package was used to convert the Seurat object to a Monocle object. Differential expression testing across the single-cell trajectory was performed using graph_test() and filtered using morgans test statistic > 0.01 and adjusted p-value < 0.05.

### CUT&RUN

CUT&RUN was performed using the CUT&RUN Assay Kit (CST) according to manufacturer’s instructions. Live sorted 60-200k mesothelial and epithelial cells were used to profile H3K27ac and 500K epithelial cells were used to profile the IgG negative control. DNA library prep was performed using the KAPA HyperPrep Kit (Roche) and sequenced on the NovaSeq 6000 with 150 bp Paired End Reads.

FastQC was used to perform quality control metrics on sequencing samples. Tru-Seq Adapters and Poly G tails were trimmed using Cutadapt (version 4.8). Alignment to the mm10 genome was performed using Bowtie2 (version 2.5.3) using the following parameters: --end-to-end --very-sensitive --no-mixed --no-discordant --phred33 -I 10 -X 700. Peaks were identified using Macs2(version 2.2.9.1) using the following parameters: --broad --broad-cutoff 0.1 --keep-dup all. To generate bigwig files for IGV visualization, deepTools (version 3.5.5) was used. bamCoverage was used with the following parameters : --binSize 20 --smoothLength 60 -- extendReads --normalizeUsing CPM --centerReads. Peak visualization was performed using deepTools (version 3.5.5) computematrix and plotHeatmap commands.

Diffbind (version 3.6.5) was used to identify differentially enriched peaks between mesothelial and epithelial cells using default parameters (summits=500)^68^. Data was normalized using edgeR. Motif analysis was performed using Homer in differentially enriched peaks (bedtools v2.31.1) peaks using findMotifsGenome.pl and -size 500.

### Quantification and Statistical Analysis

Statistical tests (2 tailed unpaired Students T-Test or One-way ANOVA with multiple comparisons) are indicated in the figure legends. Statistical significance was performed using Graphpad Prism (10.2.3). P values less than 0.05 were considered significant.

### Qupath

For ACTA2, TAGLN and ELASTIN quantification, images were analyzed with Qupath (Version 0.4.3). Control and mutant slides were stained and imaged in parallel. 3 evenly spaced lung sections were stained per animal and images were photographed using the Olympus VS200 Slide Scanner. Periphery annotations were drawn manually or defined using the tdTom reporter and TRITC pixel classifier. To quantify the percentage of ACTA2 or TAGLN positive pixels, a FITC pixel classifier of 10,000 was applied to the periphery annotation.

## Acknowledgements

We would like to thank members of the Xin Sun Lab for thoughtful discussion. We would like to thank Dr. Ben Emery and Dr. Wei Shi for sharing mouse strains. This work was supported by NIH F31HL160204 (to G.L.) and NIH R01HL172027 (to X.S). Confocal Microscopy was performed at the UCSD School of Medicine Microscopy Core (NINDS P30NS047101). Flow Cytometry was performed at the San Diego Center for Aids Research (P30 AI036214). This publication includes data generated at the UC San Diego IGM Genomics Center utilizing an Illumina NovaSeq 6000 that was purchased with funding from a National Institutes of Health SIG grant (#S10 OD026929).

## Supplemental Information

**Figure S1.**
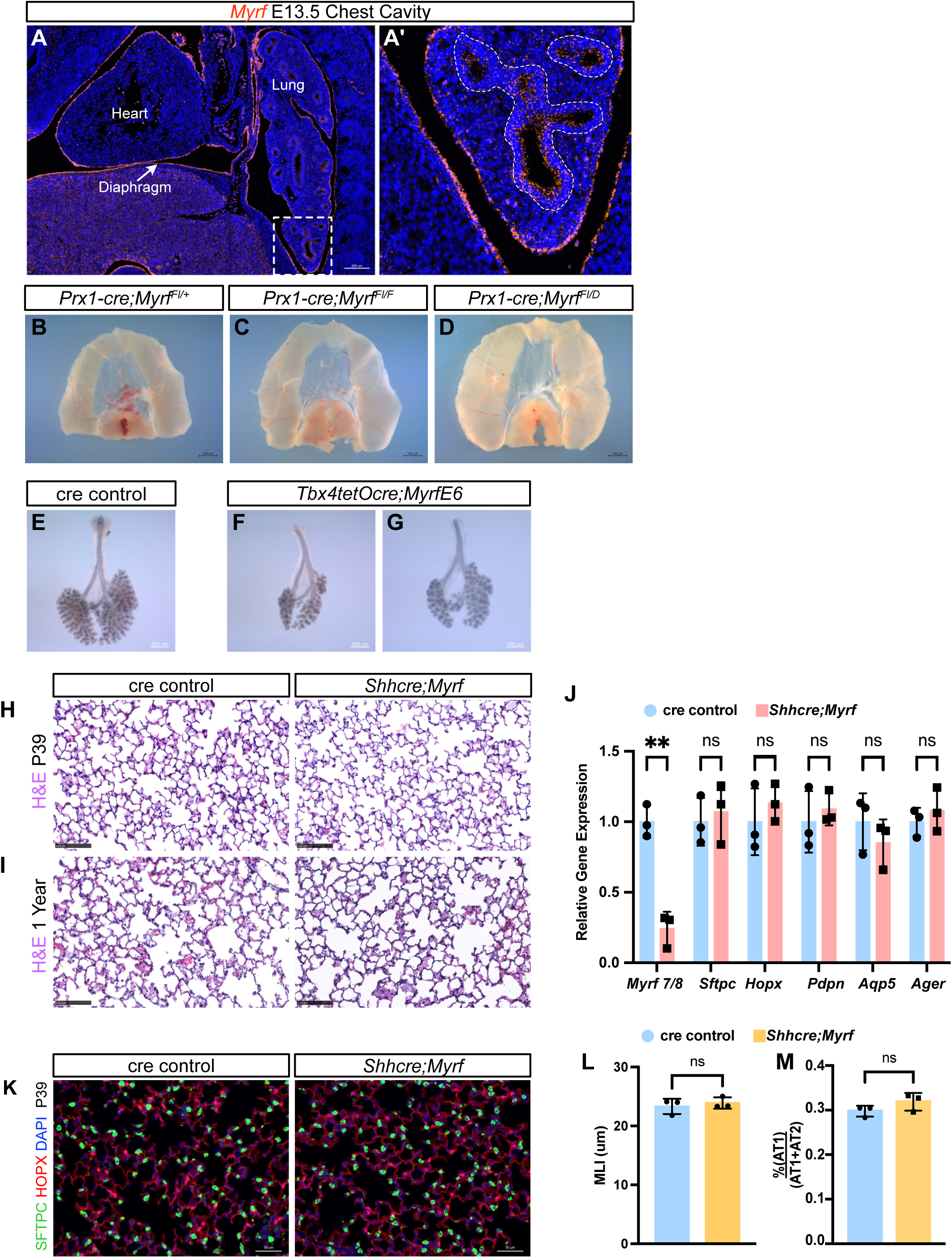
**(A)** RNAscope staining for *Myrf*(red) in a E13.5 embryonic mouse chest cavity. **(B)** Representative diaphragm image of the P42 *Prx1-cre;Myrf^Fl/+^* control. **(C)** Representative diaphragm image of the P42 *Prx1-cre;Myrf^Fl/Fl^* mutant. **(D)** Representative diaphragm image of the P42 *Prx1-Cre;Myrf^Fl/D^* mutant. **(E)** E-CADHERIN whole lung immunostaining of a representative E13.5 control lung**. (F)** E-CADHERIN whole lung immunostaining of a representative *Tbx4tetOcre;MyrfE6* mutant lung. **(G)** E-CADHERIN whole lung immunostaining of a representative *Tbx4tetOcre;MyrfE6* mutant lung. **(H)** H&E Images in P39 *Shhcre;Myrf* mutants and cre controls. **(I)** H&E Images in 1 year aged *Shhcre;Myrf* mutants and cre controls. **(J)** qRT-PCR quantification of *Myrf* and additional AT1/AT2 markers in *Shhcre;Myrf* mutants and cre controls. **(K)** Immunostaining for HOPX(red) and SPC(green) in P39 *Shhcre;Myrf* mutants and cre controls. **(L)** MLI quantification in P39 *Shhcre;Myrf* mutants and cre controls. **(M)** AT1 quantification in P39 *Shhcre;Myrf* mutants and cre controls. Students T-Test was used for S1J, S1L and S1M. ns (not significant) for P ≥ 0.05, * for P < 0.05, ** for P < 0.01, *** for P < 0.001, and **** for P < 0.0001.

**Figure S2.**
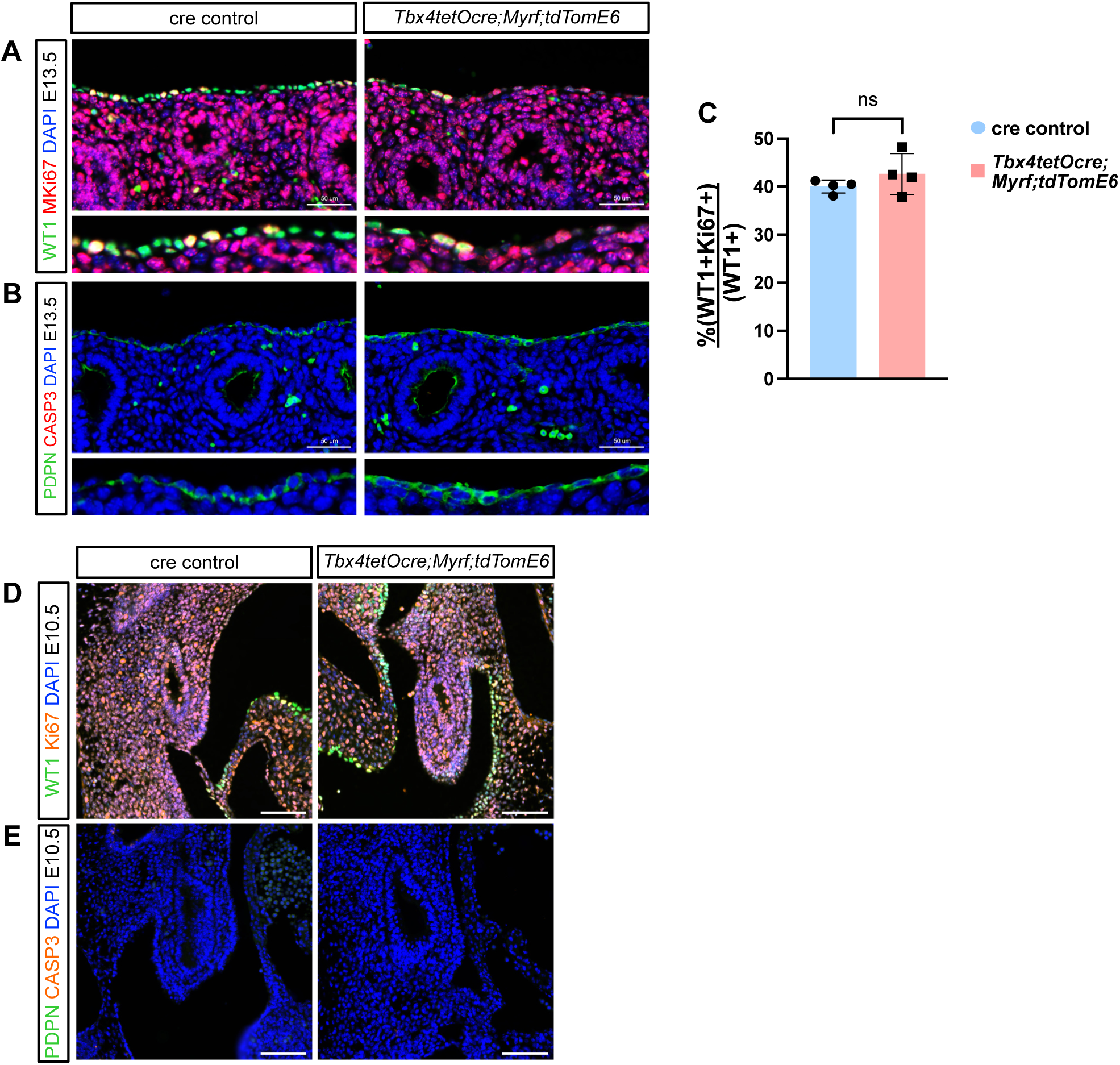
**(A)** Immunostaining for WT1(green) and MKi67(red) in the E13.5 *Tbx4tetOcre;MyrfE6* mutant and cre control. **(B)** Immunostaining for PDPN(Green) and Cleaved Caspase-3 (red) in the E13.5 *Tbx4tetOcre;MyrfE6* mutant and cre control. **(C)** Percentage of WT1+Mki67+ Cells in the in the E13.5 *Tbx4tetOcre;MyrfE6* mutant and cre control. **(D)** Immunostaining for WT1(green) and mKi67(red) in the E10.5 *Tbx4tetOcre;MyrfE6* mutant and cre control. **(E)** Immunostaining PDPN(Green) and Cleaved Caspase-3(red)in the E10.5 *Tbx4tetOcre;MyrfE6* mutant and cre control. Students T-Test was used for S2C. ns (not significant) for P ≥ 0.05, * for P < 0.05, ** for P < 0.01, and *** for P < 0.001.

**Figure S3.**
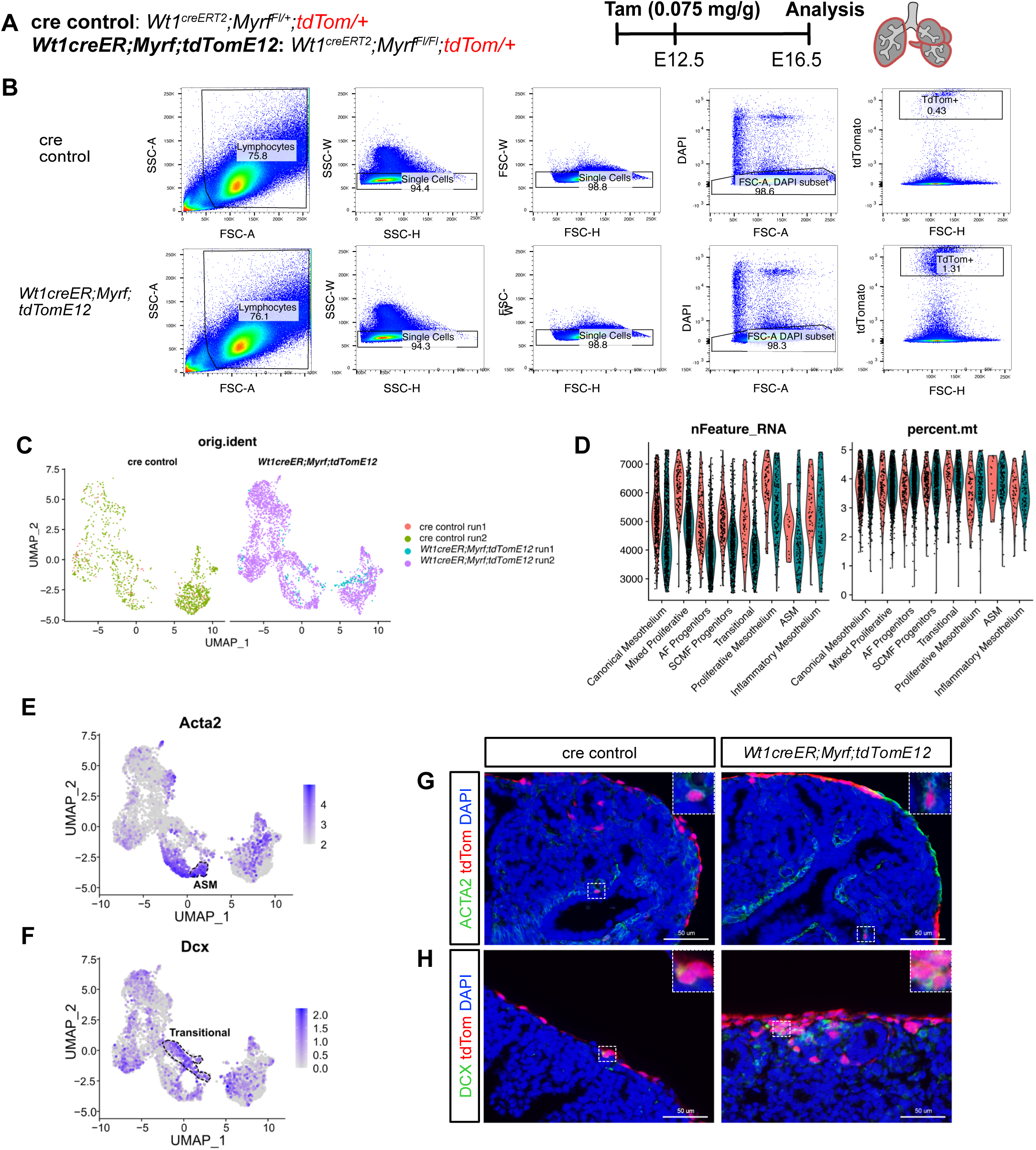
**(A)** Timeline of inactivation and analysis of the *Wt1creER;Myrf;tdTomE12* mutant and cre control. **(B)** Flow cytometry of tdTom+ lineaged traced *Wt1creER;Myrf;tdTomE12* mutants and cre controls. **(C)** Orig.ident of the 10x single cell runs. **(D)** RNA features and percent.mt of the integrated populations. **(E)** ACTA2 Featureplot of the integrated UMAP. **(F)** DCX Featureplot of the integrated UMAP. **(G)** Immunostaining for ACTA2(green) and tdTom(red). **(H)** Immunostaining for DCX(green) and tdTom (red).

**Figure S4.**
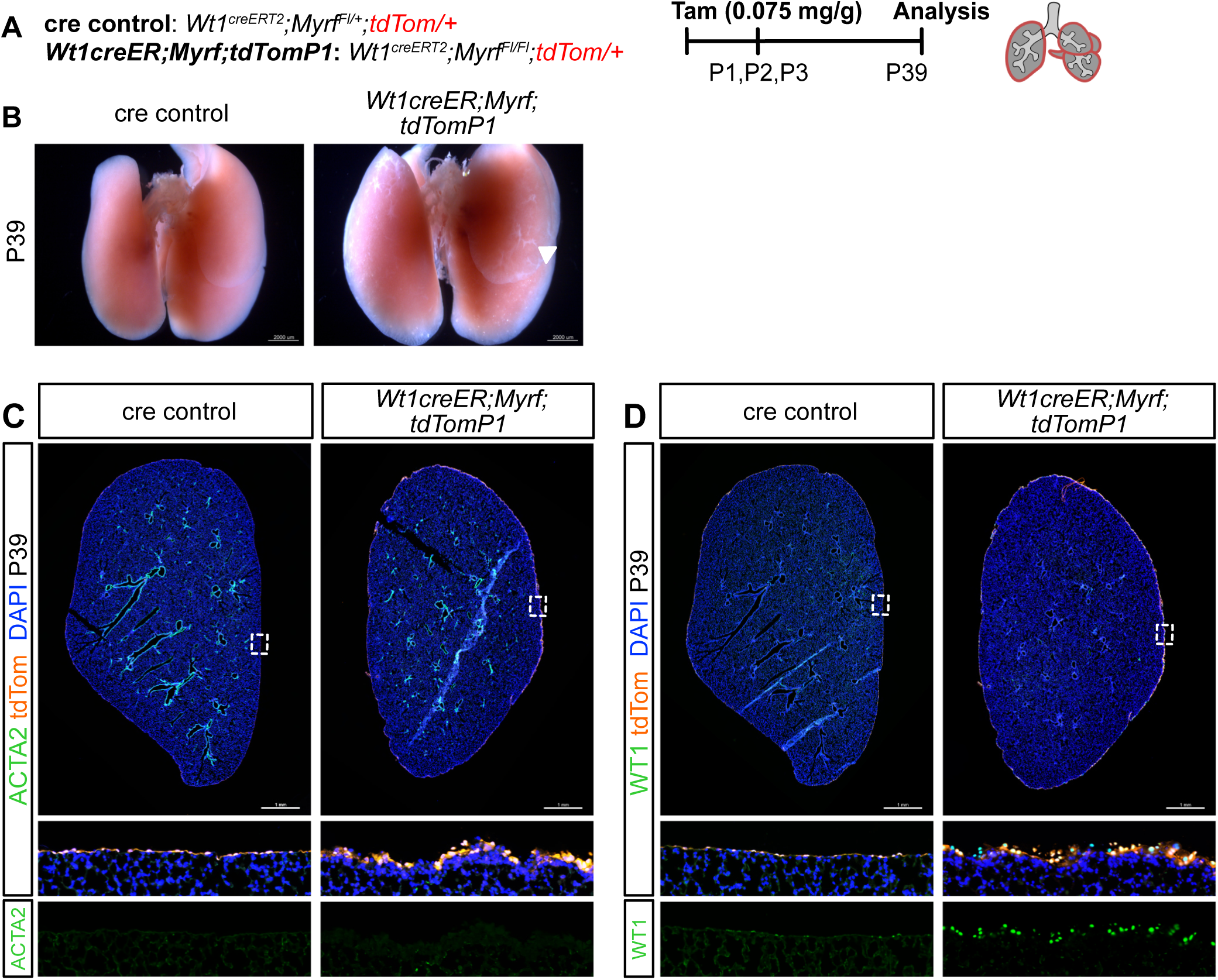
**(A)** Timeline of inactivation and analysis of the *Wt1creER;Myrf;tdTomP1* mutant and cre control. **(B)** Whole mount image of the *Wt1creER;Myrf;tdTom* mutant and cre control. **(C)** Immunostaining for ACTA2(green) and tdTom(red) in the*Wt1creER;Myrf;tdTomP1* mutant and cre control. **(D)** Immunostaining for WT1(green) and tdTom(red) in the *Wt1creER;Myrf;tdTomP1* mutant and cre control.

**Figure S5.**
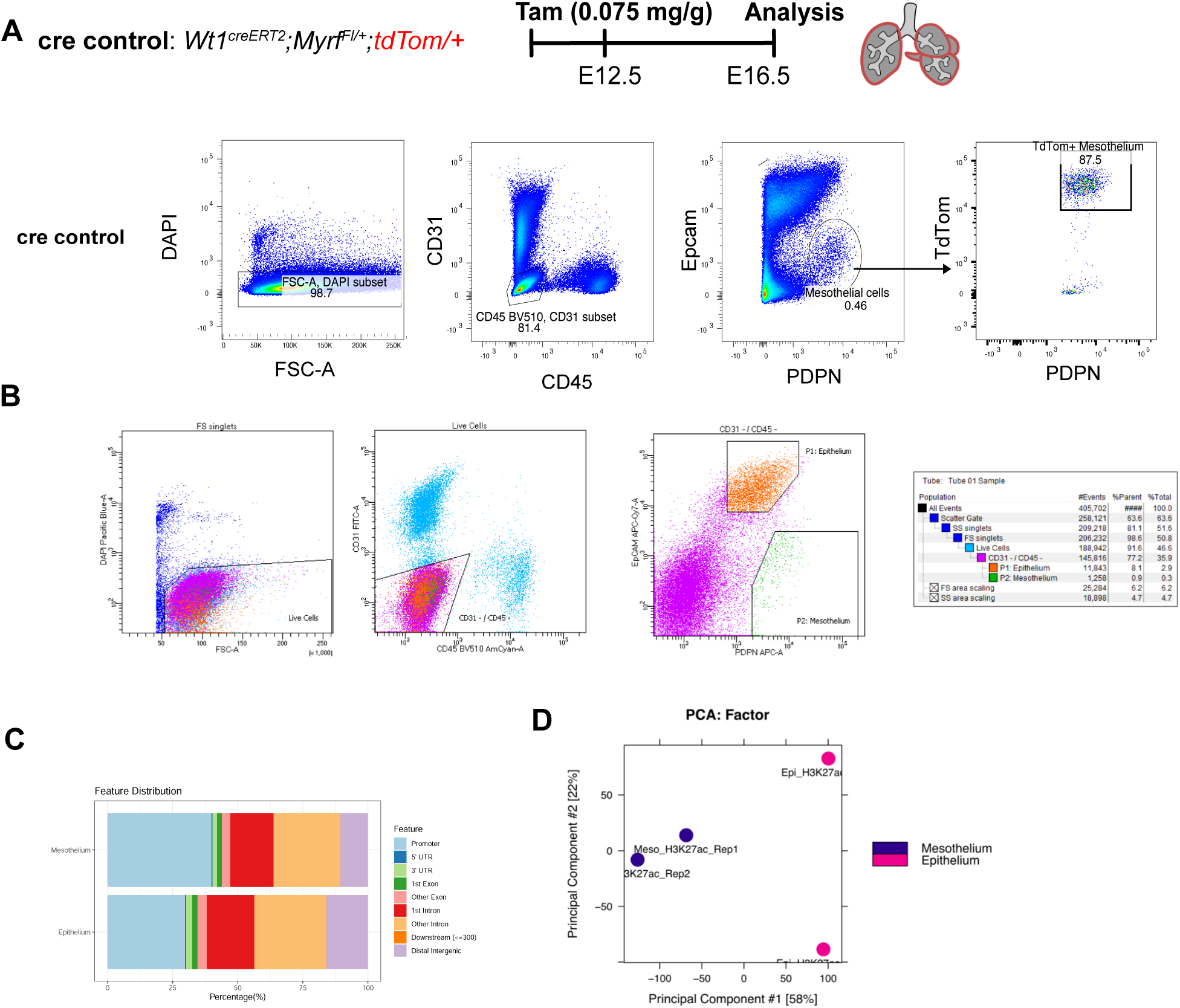
**(A)** Validation of flow cytometry panel using *Wt1creER;Myrf;tdTomE12* control lungs. tdTom+ Mesothelial cells are CD31-CD45-EPCAM-PDPN+. Epithelial cells are CD31-CD45- EPCAM+PDPN+. **(B)** Sample flow cytometry gating used to sort mesothelium and epithelium populations. **(C)** Feature Distribution of intersected H3K27ac peaks in the mesothelium and epithelium. **(D)** Diffbind PCA plot.

**Figure S6.**
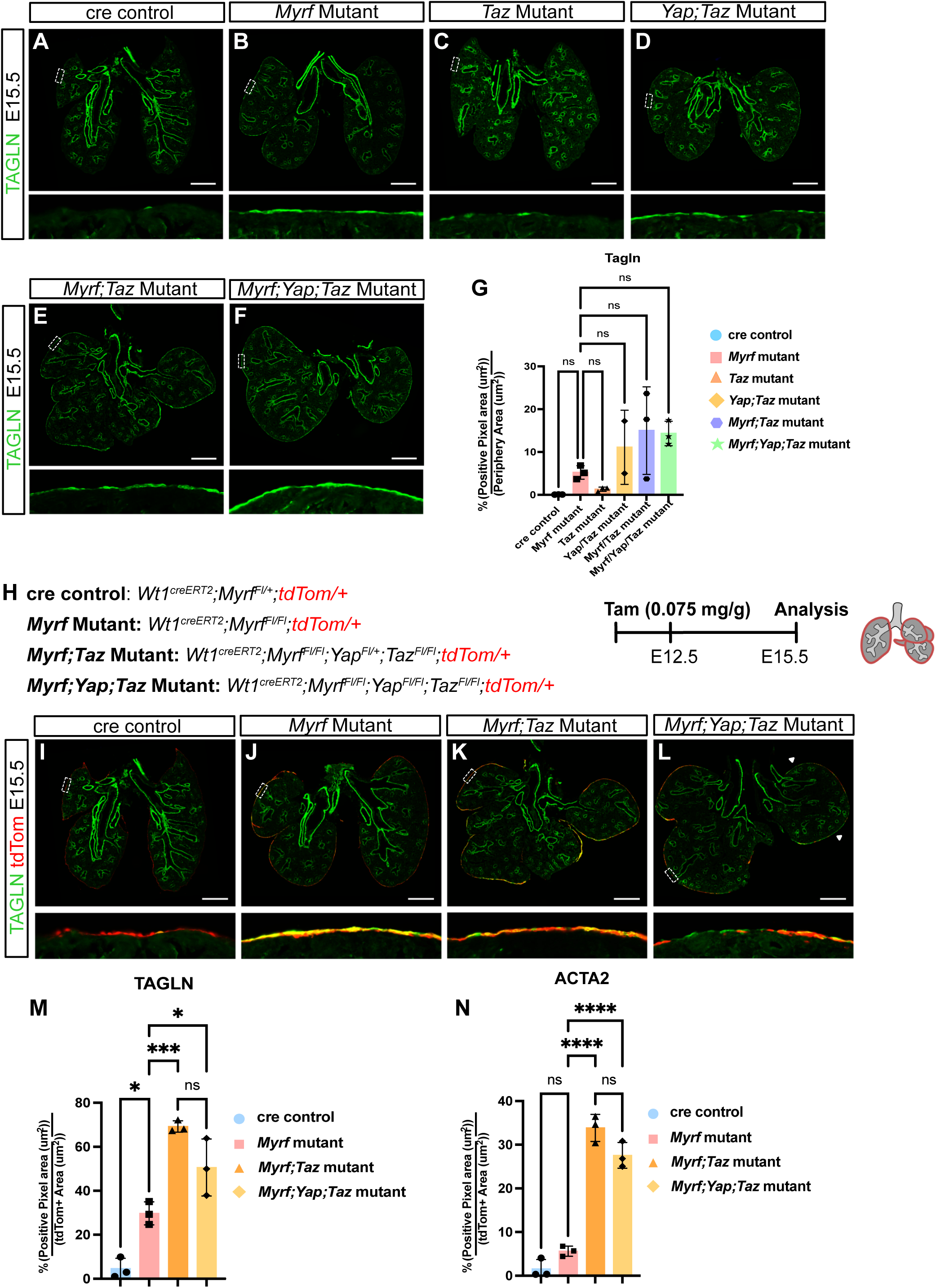
**(A)** Representative TAGLN(green) immunostaining of the cre control. **(B)** Representative TAGLN(green) immunostaining of the *Myrf* mutant. **(C)** Representative TAGLN(green) immunostaining of the *Taz* mutant. **(D)** Representative TAGLN(green) immunostaining of the *Yap;Taz* mutant. **(E)** Representative TAGLN(green) immunostaining of the *Myrf;Taz* mutant. **(F)** Representative TAGLN(green) immunostaining of the *Myrf;Yap;Taz* mutant. **(G)** Quantification of TAGLN positive pixel area over the periphery area in the cre control, *Myrf*, *Taz, Yap;Taz, Myrf;Taz*, and *Myrf;Yap;Taz* mutants. *Myrf* mutant interactions are displayed in the graph. **(H)** Timeline of inactivation and analysis of the cre control, *Myrf*, *Myrf;Taz*, and *Myrf;Yap;Taz* mutants crossed to a tdTom reporter. **(I)** Representative TAGLN(green) and tdTom(red) immunostaining of the cre control. **(J)** Representative TAGLN(green) and tdTom(red) immunostaining of the *Myrf* mutant. **(K)** Representative TAGLN(green) and tdTom(red) immunostaining of the *Myrf;Taz* mutant. **(L)** Representative TAGLN(green) and tdTom(red) immunostaining of the *Myrf;Yap;Taz* mutant. **(M)** Quantification of TAGLN positive pixel area over the tdTom area in the cre control, *Myrf*, *Myrf;Taz*, and *Myrf;Yap;Taz* mutants. **(N)** Quantification of ACTA2 positive pixel area over the tdTom area in the cre control, *Myrf*, *Myrf;Taz*, and *Myrf;Yap;Taz* mutants.

**Figure S7.**
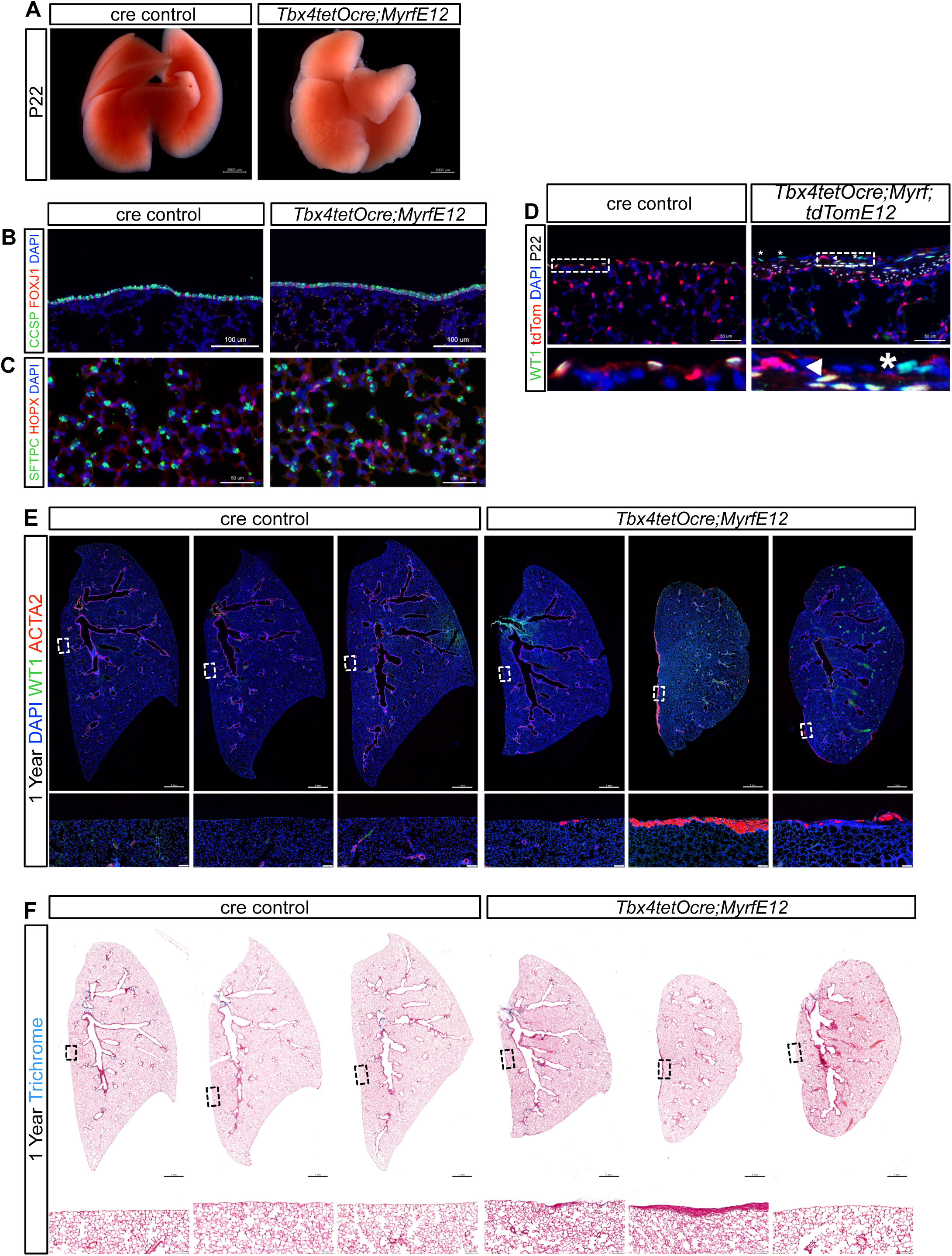
**(A)** Representative dorsal whole mount images of the *Tbx4tetOcre;MyrfE12* mutant and cre control at P22. **(B)** Immunostaining for CCSP(green) and FOXJ1(red). **(C)** Immunostaining for HOPX(red) and SFTPC(green). **(D)** Immunostaining for WT1(green) and tdTom(red) in the P22 *Tbx4tetOcre;Myrf;tdTomE12* mutant and cre control. **(E)** Immunostaining for WT1(green) and ACTA2(red) in 1-year aged *Tbx4tetOcre;MyrfE12* mutants and cre controls. **(F)** Representative trichrome stains of 1-year aged *Tbx4tetOcre;MyrfE12* mutants and cre controls.

TableS1.xls E13.5 *Tbx4tetOcre;MyrfE6* Bulk RNA-Seq

TableS2.xls E16.5 scRNA-seq population markers

TableS3.xls E16.5 *Wt1creER;Myrf;tdTom;E12* canonical mesothelium DE gene

TableS4.xls Monocle 3 Graph auto-correlation analysis

TableS5.xls Diffbind peak characterization TableS6.xls qPCR Primers

## Notes

### Competing Interest Statement

The authors have declared no competing interest.

