## Supplemental Figures for "MYRF is Essential in Mesothelial Cells to Promote Lung Development and Maturation"

**Figure S1**

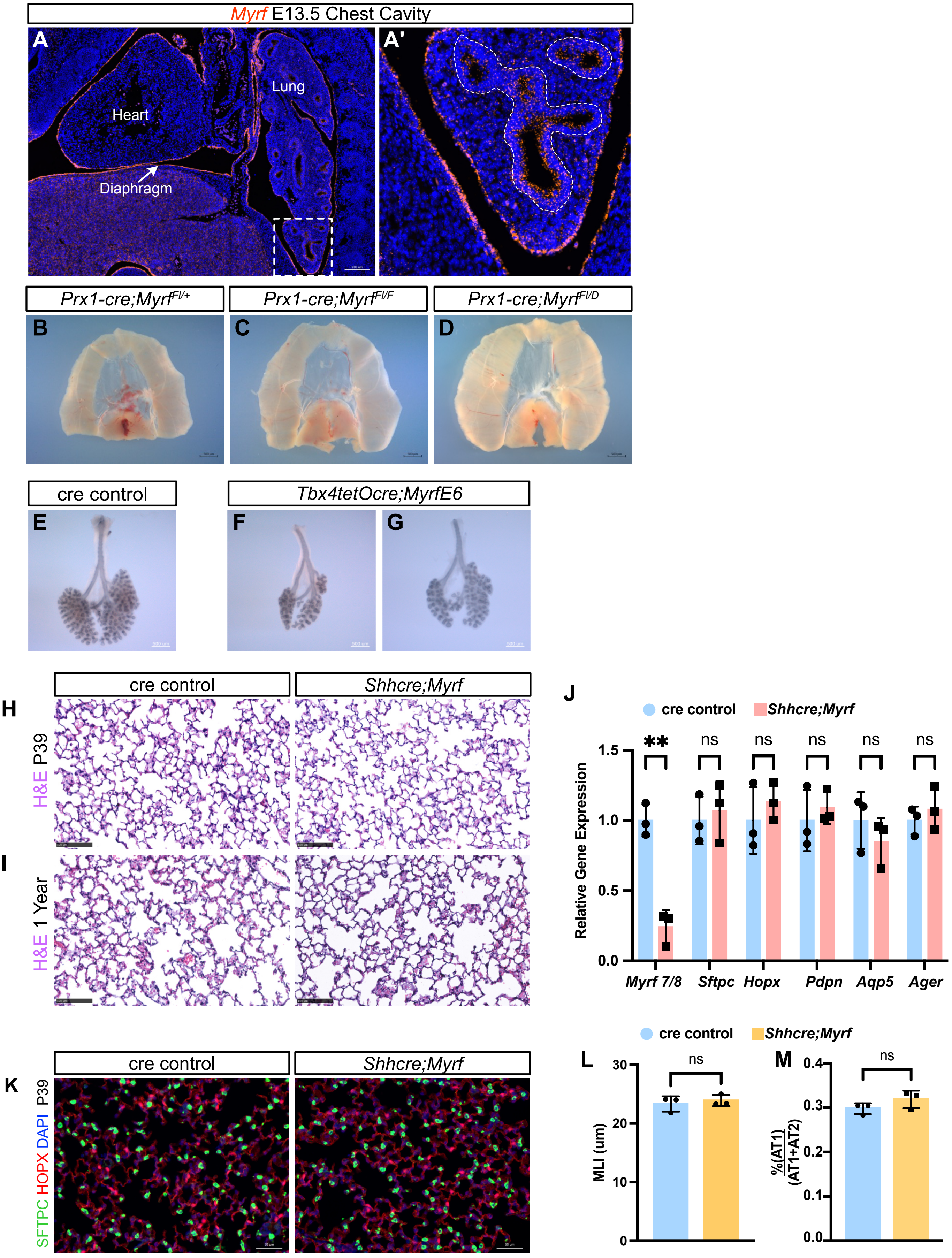

Myrf E13.5 Chest Cavity
A
A'
Heart
Lung
Diaphragm
Prx1-cre;Myrf^Fl/+
Prx1-cre;Myrf^Fl/F
Prx1-cre;Myrf^Fl/D
B
C
D
cre control
Tbx4tetOcre;MyrfE6
E
F
G
cre control
Shhcre;Myrf
H
H&E P39
I
H&E 1 Year
J
cre control
Shhcre;Myrf
Relative Gene Expression
1.5
1.0
0.5
0.0
**
ns
ns
ns
ns
ns
Myrf 7/8
Sftpc
Hopx
Pdpn
Aqp5
Ager
K
cre control
Shhcre;Myrf
SFTPC HOPX DAPI P39
cre control
Shhcre;Myrf
L
MLI (um)
30
20
10
0
ns
M
%(AT1)/(AT1+AT2)
0.4
0.3
0.2
0.1
0.0
ns

Figure S2

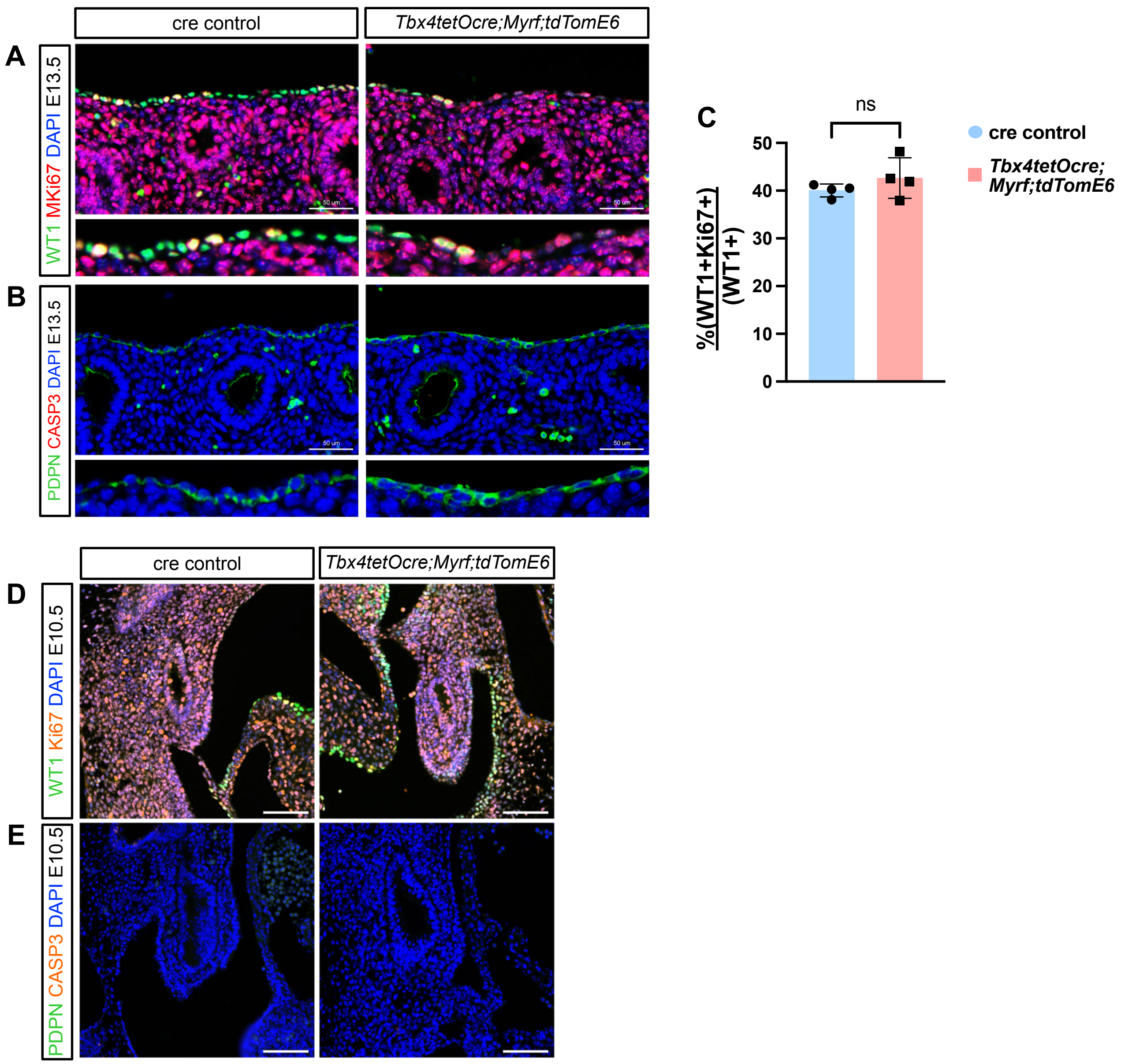

cre control
Tbx4tetOcre;Myrf;tdTomE6
A
WT1 MKi67 DAPI E13.5
B
PDPN CASP3 DAPI E13.5
C
ns
%(WT1+Ki67+)/(WT1+)
50
40
30
20
10
0
cre control
Tbx4tetOcre; Myrf;tdTomE6
D
WT1 Ki67 DAPI E10.5
E
PDPN CASP3 DAPI E10.5

### Figure S3

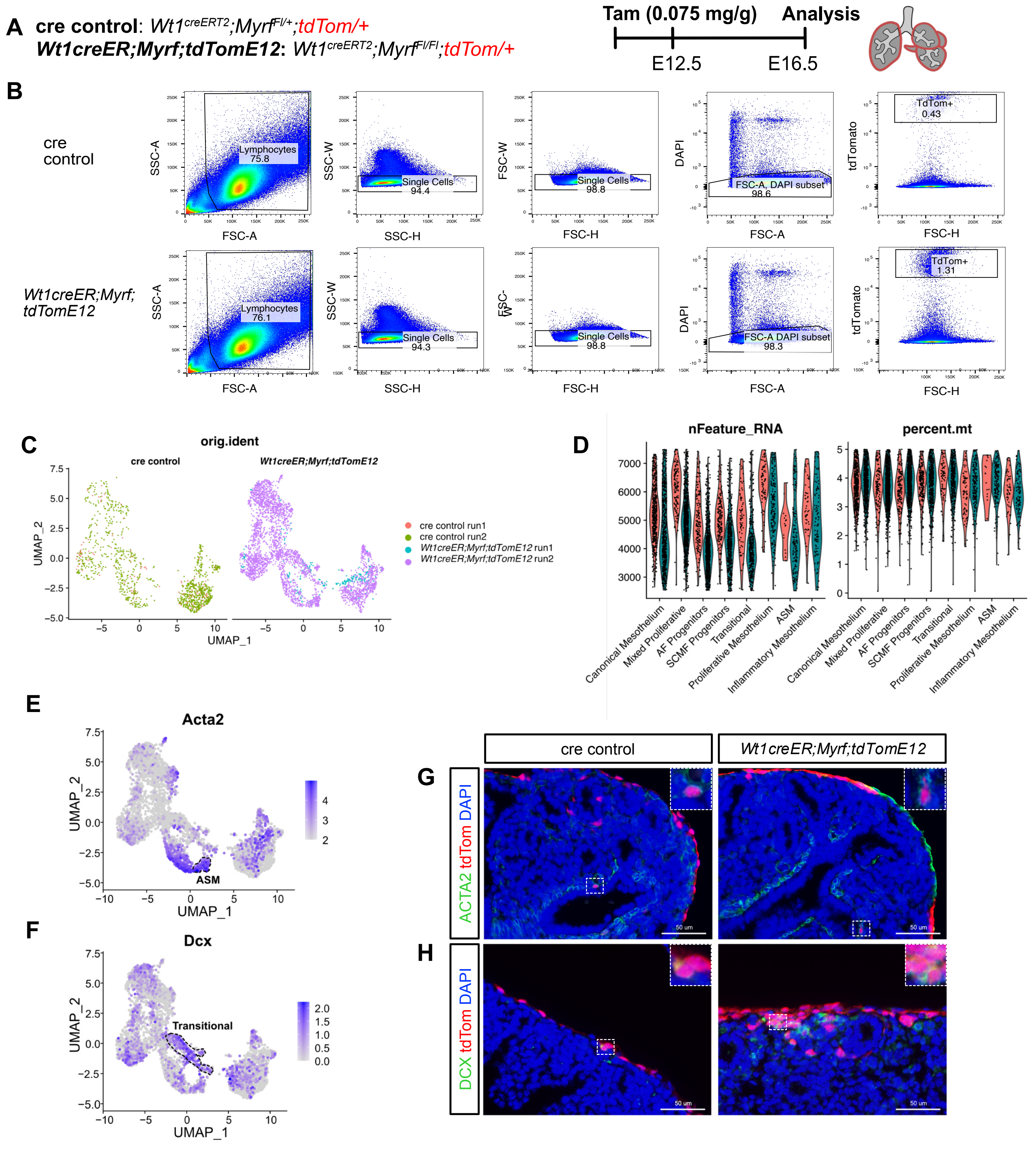
A
cre control: Wt1creERT2;MyrfFl/+;tdTom/+
Wt1creER;Myrf;tdTomE12: Wt1creERT2;MyrfFl/Fl;tdTom/+
Tam (0.075 mg/g)
Analysis
E12.5
E16.5
B
cre control
Wt1creER;Myrf; tdTomE12
Lymphocytes 75.8
Single Cells 94.4
Single Cells 98.8
FSC-A, DAPI subset 98.6
TdTom+ 0.43
Lymphocytes 76.1
Single Cells 94.3
Single Cells 98.8
FSC-A DAPI subset 98.3
TdTom+ 1.31
SSC-A
SSC-W
FSC-W
DAPI
tdTomato
FSC-A
SSC-H
FSC-H
C
orig.ident
cre control
Wt1creER;Myrf;tdTomE12
UMAP_2
UMAP_1
cre control run1
cre control run2
Wt1creER;Myrf;tdTomE12 run1
Wt1creER;Myrf;tdTomE12 run2
D
nFeature_RNA
percent.mt
Canonical Mesothelium
Mixed Proliferative
AF Progenitors
SCMF Progenitors
Transitional
Proliferative Mesothelium
ASM
Inflammatory Mesothelium
E
Acta2
ASM
F
Dcx
Transitional
cre control
Wt1creER;Myrf;tdTomE12
G
ACTA2 tdTom DAPI
H
DCX tdTom DAPI
50 um

**Figure S4**

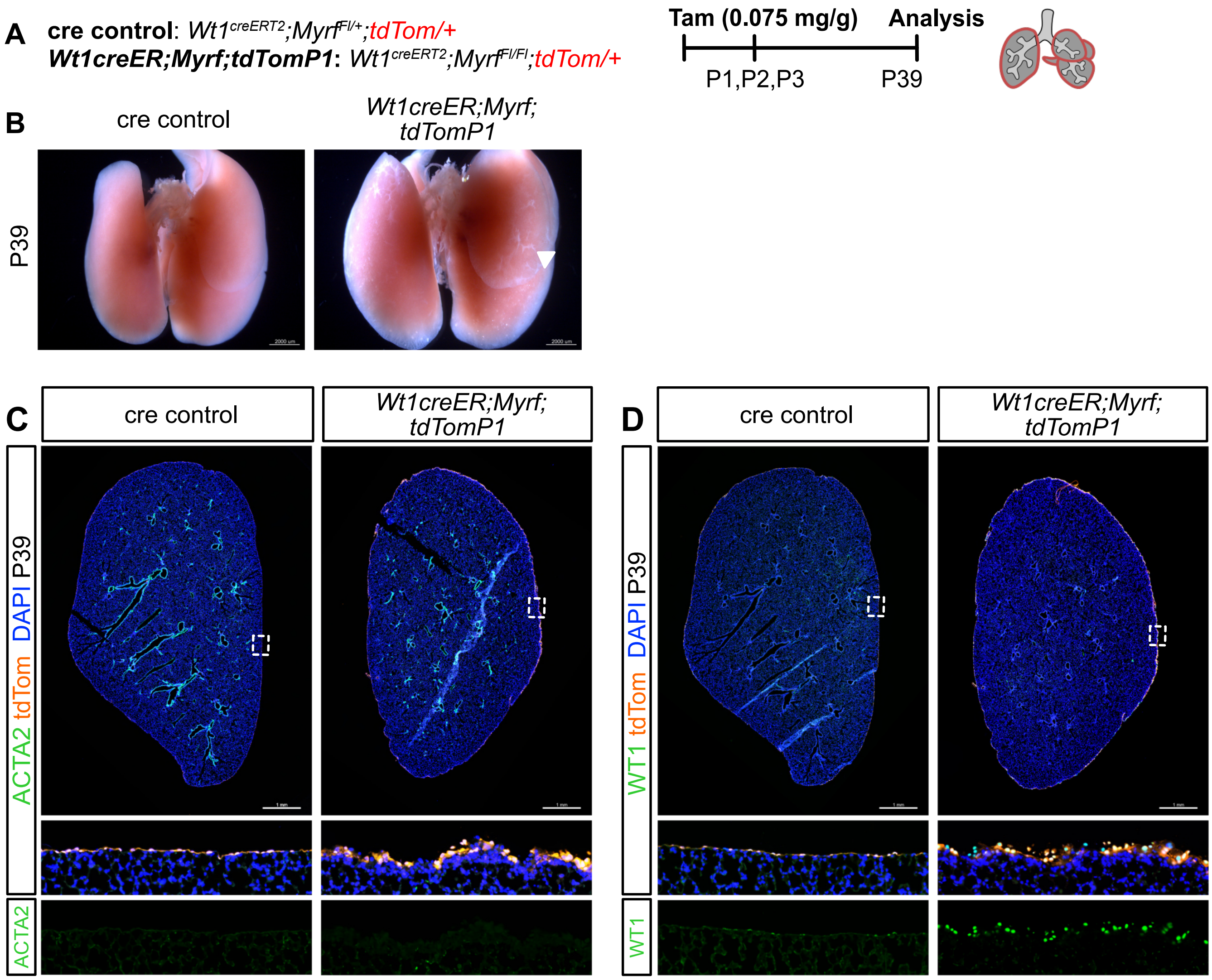
A
cre control: Wt1creERT2;MyrfFl/+;tdTom/+
Wt1creER;Myrf;tdTomP1: Wt1creERT2;MyrfFl/Fl;tdTom/+
Tam (0.075 mg/g)
Analysis
P1,P2,P3
P39
B
cre control
Wt1creER;Myrf; tdTomP1
P39
C
cre control
Wt1creER;Myrf; tdTomP1
ACTA2 tdTom DAPI P39
ACTA2
D
cre control
Wt1creER;Myrf; tdTomP1
WT1 tdTom DAPI P39
WT1

### Figure S5

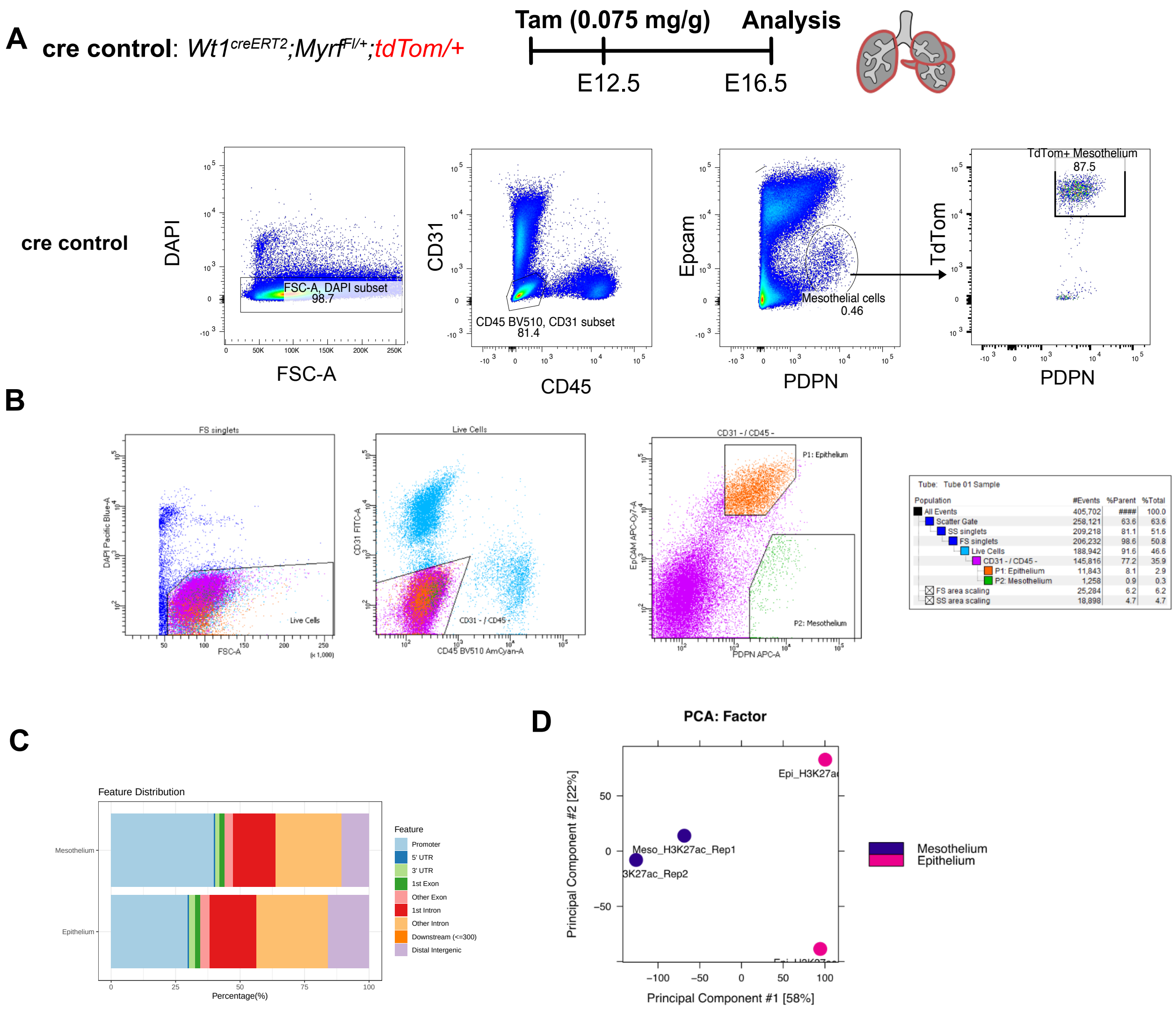


| Population | #Events | %Parent | %Total |
|---|---|---|---|
| All Events | 405,702 | #### | 100.0 |
| Scatter Gate | 258,121 | 63.6 | 63.6 |
| SS singlets | 209,218 | 81.1 | 51.6 |
| FS singlets | 206,232 | 98.6 | 50.8 |
| Live Cells | 188,942 | 91.6 | 46.6 |
| CD31 - / CD45 - | 145,816 | 77.2 | 35.9 |
| P1: Epithelium | 11,843 | 8.1 | 2.9 |
| P2: Mesothelium | 1,258 | 0.9 | 0.3 |
| FS area scaling | 25,284 | 6.2 | 6.2 |
| SS area scaling | 18,898 | 4.7 | 4.7 |

**Figure S6**

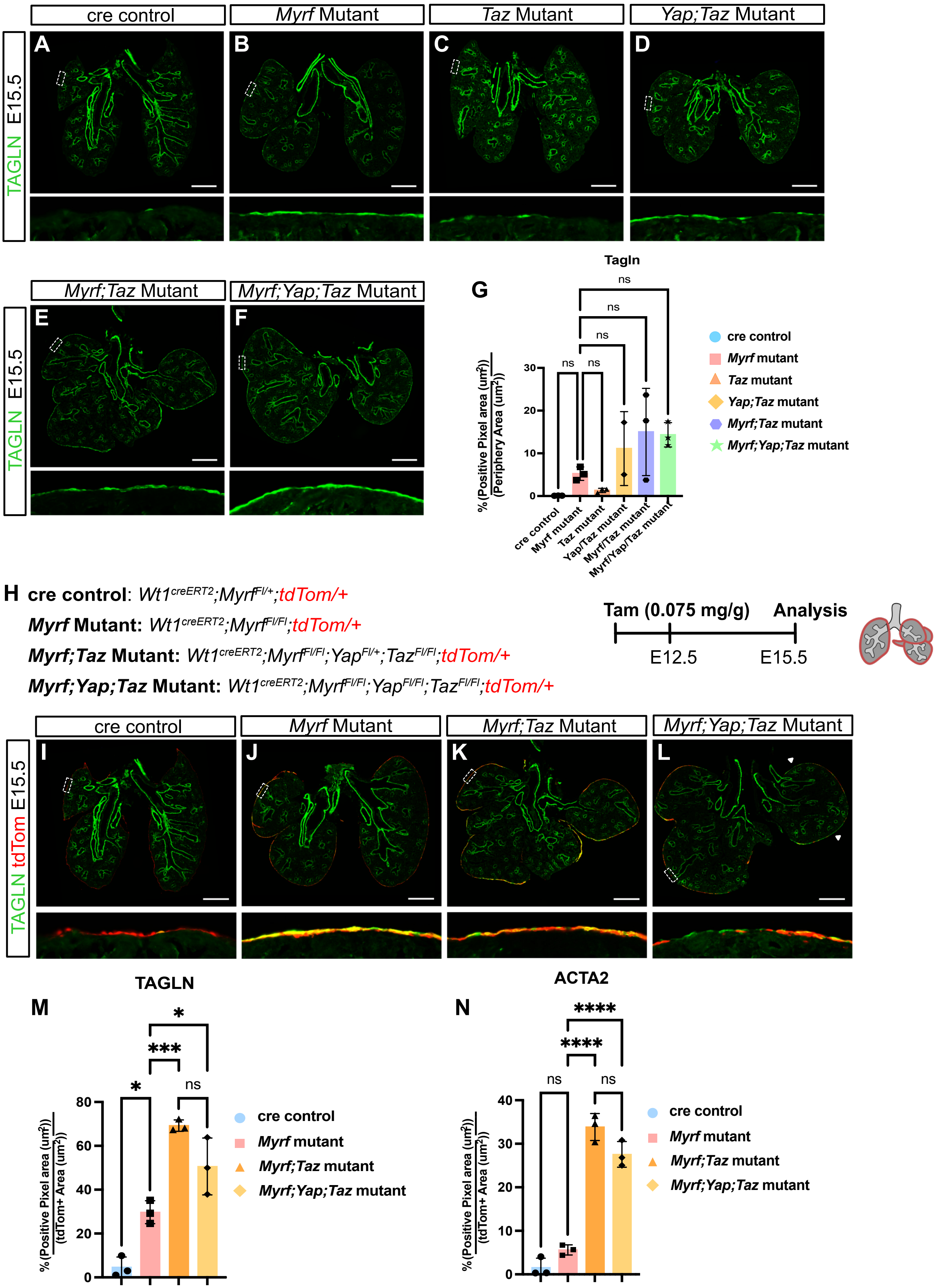
cre control
Myrf Mutant
Taz Mutant
Yap;Taz Mutant
TAGLN E15.5
A
B
C
D
Myrf;Taz Mutant
Myrf;Yap;Taz Mutant
E
F
G
Tagln
ns
ns
ns
ns
ns
% (Positive Pixel area (um²) / (Periphery Area (um²))
30
20
10
0
cre control
Myrf mutant
Taz mutant
Yap/Taz mutant
Myrf/Taz mutant
Myrf/Yap/Taz mutant
cre control
Myrf mutant
Taz mutant
Yap;Taz mutant
Myrf;Taz mutant
Myrf;Yap;Taz mutant
H
cre control: Wt1creERT2;MyrfFl/+;tdTom/+
Myrf Mutant: Wt1creERT2;MyrfFl/Fl;tdTom/+
Myrf;Taz Mutant: Wt1creERT2;MyrfFl/Fl;YapFl/+;TazFl/Fl;tdTom/+
Myrf;Yap;Taz Mutant: Wt1creERT2;MyrfFl/Fl;YapFl/Fl;TazFl/Fl;tdTom/+
Tam (0.075 mg/g)
Analysis
E12.5
E15.5
cre control
Myrf Mutant
Myrf;Taz Mutant
Myrf;Yap;Taz Mutant
TAGLN tdTom E15.5
I
J
K
L
M
TAGLN
*
***
*
ns
% (Positive Pixel area (um²) / (tdTom+ Area (um²))
80
60
40
20
0
cre control
Myrf mutant
Myrf;Taz mutant
Myrf;Yap;Taz mutant
N
ACTA2
****
****
ns
ns
% (Positive Pixel area (um²) / (tdTom+ Area (um²))
40
30
20
10
0
cre control
Myrf mutant
Myrf;Taz mutant
Myrf;Yap;Taz mutant

**Figure S7**

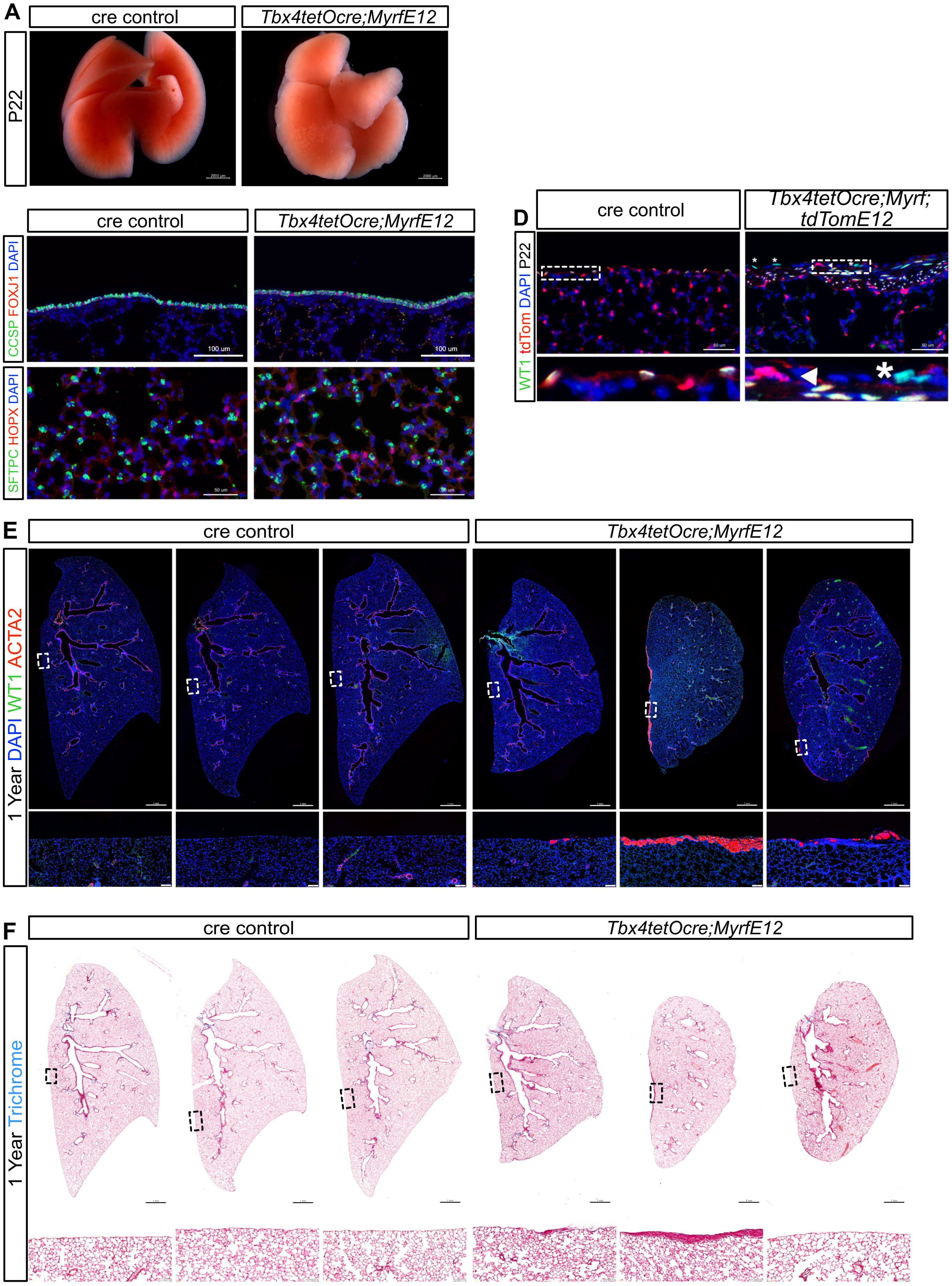
A
cre control
Tbx4tetOcre;MyrfE12
P22
B
cre control
Tbx4tetOcre;MyrfE12
CCSP FOXJ1 DAPI
100 um
C
SFTPC HOPX DAPI
50 um
D
cre control
Tbx4tetOcre;Myrf; tdTomE12
WT1 tdTom DAPI P22
E
cre control
Tbx4tetOcre;MyrfE12
1 Year DAPI WT1 ACTA2
F
cre control
Tbx4tetOcre;MyrfE12
1 Year Trichrome
